# Rothmund-Thomson Syndrome-like RECQL4 truncating mutations cause a haploinsufficient low bone mass phenotype in mice

**DOI:** 10.1101/2020.11.11.379214

**Authors:** Wilson Castillo-Tandazo, Ann E Frazier, Natalie A Sims, Monique F Smeets, Carl R Walkley

## Abstract

Rothmund-Thomson Syndrome (RTS) is an autosomal recessive disorder characterized by poikiloderma, sparse or absent hair, and defects in the skeletal system such as bone hypoplasia, short stature, low bone mass, and an increased incidence of osteosarcoma. RTS type 2 patients typically present with germline compound bi-allelic protein-truncating mutations of *RECQL4*. As existing murine models predominantly employ *Recql4* null alleles, we have here attempted to more accurately model the mutational spectrum of RTS by generating mice with patient-mimicking truncating *Recql4* mutations. We found that truncating mutations impaired stability and subcellular localization of RECQL4, which translated to a homozygous embryonic lethality and haploinsufficient low bone mass and reduced cortical bone thickness phenotypes. Combination of a truncating mutation with a conditional *Recql4* null allele demonstrated that these defects were intrinsic to the osteoblast lineage. However, the truncating mutations did not promote tumorigenesis, even after exposure to irradiation. We also utilized murine *Recql4* null cells to assess the impact of a wider range of human *RECQL4* mutations using an *in vitro* complementation assay. We found differential effects of distinct RECQL4 mutations. While some created unstable protein products, others altered subcellular localization of the protein. Interestingly, the severity of the phenotypes correlated with the extent of protein truncation. Collectively, our results reveal that truncating RECQL4 mutations lead to the development of an osteoporosis-like phenotype through defects in early osteoblast progenitors in mice and identify RECQL4 gene dosage as a novel regulator of bone mass.

## Introduction

Rothmund-Thomson syndrome (RTS) (OMIM #268400) is a rare autosomal recessive disorder that presents with skin rash (poikiloderma; areas of hypopigmentation, hyperpigmentation, telangiectasias and atrophy of the skin), sparse or absent hair, juvenile cataracts, gastrointestinal and skeletal complications (1, 2). Approximately 75% of patients have skeletal abnormalities, including bone hypoplasia, short stature, polydactyly, and low bone mass (3). Furthermore, this syndrome is frequently associated with osteosarcoma (bone cancer) and other malignancies (1, 2, 4, 5). RTS is classified into two forms: RTS type 1, where patients present with juvenile cataracts and have frequent mutations of *ANAPC1,* but do not have increased incidence of osteosarcoma (6); and RTS type 2, where the majority of patients harbor biallelic mutations of *RECQL4* and have a significantly increased incidence of osteosarcoma (2).

*RECQL4* is located on the long arm of chromosome eight (8q24.3) (7). The reported mutational spectrum includes the introduction of early stop codons, frameshift mutations, as well as deletions within numerous short introns (<100 bp) that can impair RNA splicing, resulting in protein truncations and loss-of-function alleles (8). The *RECQL4* gene encodes a protein of 1,208 amino acids (aa) that has three well-characterized domains. The N-terminal region shares sequence homology to the essential yeast DNA replication factor Sld2. In higher eukaryotes, this Sld2-like homology domain is unique to RECQL4 (9, 10). Studies have shown that this region has roles in DNA replication and DNA repair, and that it is critical for viability (11–13). The highly conserved central RecQ helicase domain contains an ATPase core with seven motifs that couple ATP hydrolysis to double-stranded DNA (dsDNA) separation (14). This ATP-dependent helicase activity was presumed to be critical for the function of RECQL4. However, using mice with a knock-in mutation (K525A) that inactivates the ATP-dependent helicase function, we recently reported that homozygous mice displayed normal embryonic development, body weight, hematopoiesis, B and T cell development, and physiological DNA damage repair (15). Finally, the C-terminal region harbors both the R4ZBD domain, in place of an RQC domain seen in other RecQ helicases, and a small C-terminal domain (CTD; 1117-1208aa) associated with DNA-binding affinity (16). Regarding intracellular localization, RECQL4 is primarily localized in the nucleus but has also been reported in the cytoplasm (17, 18). Its distribution within these compartments however, depends on the cell type and the phase of the cell cycle (17).

RTS Type 2 patients have a high incidence of skeletal abnormalities and osteosarcoma. In a clinical cohort study evaluating 41 RTS patients, 32% developed osteosarcoma (2). Furthermore, an independent cohort reported that the median age at diagnosis in RTS patients was ten years of age (19). This is significantly younger than the median age of sporadic osteosarcoma, which is sixteen years (20). Importantly, we and others have previously reported that in mice the deletion of *Recql4* in pre-osteoblasts or limb bud progenitors caused shorter bones and reduced bone volume (21, 22). However, neither model developed osteosarcoma, even in combination with *Tp53* deficiency (22). In contrast to these models that generated null alleles, RTS patients predominantly have compound heterozygous *RECQL4* mutations that are predicted to generate truncated protein products (4). More than half of these truncate the protein before or within the helicase domain and result in a substantially increased risk of developing osteosarcoma compared to non-truncating mutations (5). Therefore, it is critical to determine the *in vivo* effects of germline truncating *Recql4* mutations in normal homeostasis and tumor development.

To more faithfully model the RTS-relevant *RECQL4* mutation spectrum beyond existing null alleles, we have generated mice bearing truncating mutations that map closely to those reported in RTS patients. Here we show that these truncating mutations affected stability and subcellular localization of RECQL4, which translated to a homozygous developmental lethality and a haploinsufficient low bone mass phenotype through defects in early osteoblast progenitors. Additionally, we observed that the severity of the defect was related to the degree of the truncation, suggesting that gene dosage is an important determinant of the bone phenotype. However, unlike in RTS type 2 patients, these RECQL4 mutations were not sufficient in isolation to initiate tumorigenesis in mice, even after exposure to irradiation. This would suggest that additional molecular and cellular changes are required for the full spectrum of RTS phenotypes to develop.

## Results

### Truncating mutations of RECQL4 affect protein expression levels and cause developmental lethality in homozygotes

To understand the *in vivo* impact of truncating RECQL4 mutations, we generated two novel mouse *Recql4* alleles (15). These new mutations were similar to common mutations seen in RTS type 2 patients (Fig 1A). The p.Gly522GlufsTer43 (G522Efs) mutation, comparable to the human p.Cys525AlafsX33 (C525Afs) mutation, was created by a two-base pair insertion (c.1646_1647insGA). The frameshift caused a premature stop codon 44aa downstream, resulting in a predicted protein of 566aa lacking the majority of the helicase domain and all of the C-terminal domain. The second allele was a p.Arg347* (R347X) mutation, a nonsense mutation (c.1122C>T) identified from an N-ethyl-N-nitrosourea (ENU) mutagenesis collection, similar to p.Arg350GlyfsX21 (R350Gfs) in RTS patients. This mutation yielded a predicted 347aa protein lacking both the helicase and C-terminal domains entirely (Fig 1B). To verify the genotypes of these mice, we used PCR-restriction fragment length polymorphism (PCR-RFLP) for the G522Efs allele, and competitive allele-specific PCR (KASP) assay for the R347X allele, both of which confirmed the correct genotype (S1 Fig).

**Fig. 1.**
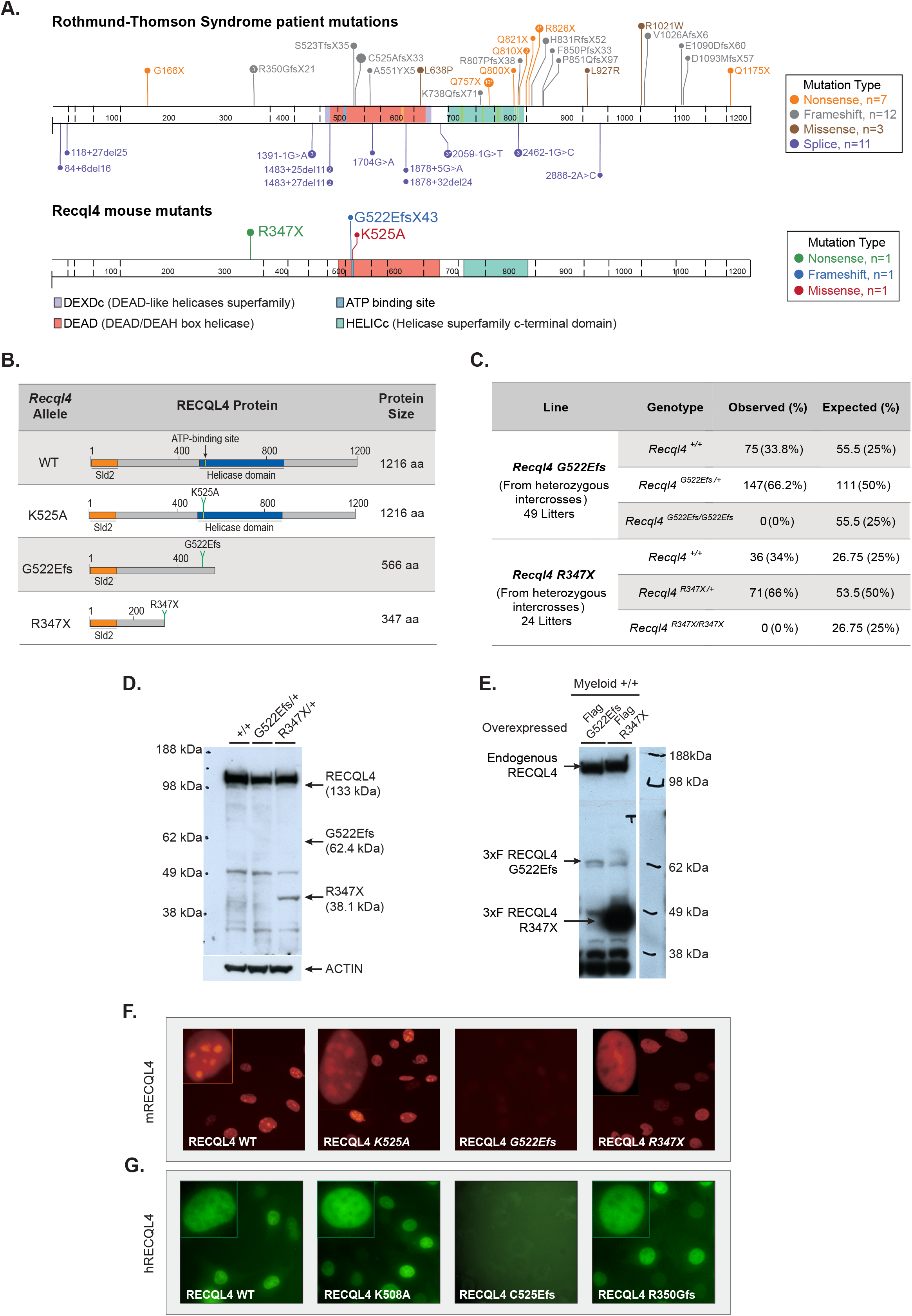
Truncating RECQL4 mutations G522Efs and R347X affect protein expression and localization differently and are homozygous embryo lethal. (A) Schematic illustration of RECQL4 mutations reported in RTS patients and murine mutations used in this study. Image generated using Protein Painter (PeCan portal, St Jude’s). (B) *Recql4* mutations and their corresponding predicted protein products. (C) Breeding data from 49 litters of *Recql4^G522Efs/+^* and 24 litters of *Recql4^R347X/+^* intercrosses. Observed and expected mendelian frequencies of the indicated genotypes are shown. No statistical significance was achieved. (D) Western blot of thymocyte lysates from *Recql4^+/+^, Recql4^G522Efs/+^,* and *Recql4^R347X/+^* probed with anti-mouse RECQL4 (clone 3B10; top). The same blot re-probed with anti-Actin (bottom). (E) Western blot of lysates from HoxB8 immortalized *R26*-CreER^T2^ *Recql4^+/+^* infected with MSCV puro 3xFlag RECQL4 and probed with anti-RECQL4 (clone 3B10; top) and M2 anti-Flag antibody (bottom). (F) Fluorescent microscopy of RECQL4 expression in Kusa 4b10 cells with murine mCherry fused WT, K525A, G522Efs, and R347X and (G) human EGFP fused WT, K508A, C525Efs, and R350Gfs mutations.

The heterozygous *Recql4^R347X/+^* and *Recql4^G522Efs/+^* mice were viable and fertile. To determine if individual homozygous truncating mutants were viable, the respective heterozygous mice were inbred. We did not recover any *Recql4^R347X/R347X^* or *Recql4^G522Efs/G522Efs^* pups at genotyping (day 7-10 after birth), indicating that the homozygous mutants were developmentally lethal (Fig 1C). We have not established the time point in development at which the respective mutants are no longer viable.

Next, we investigated the *in vivo* expression of the predicted truncated proteins. We prepared lysates from the thymus of germline heterozygous mutants of each respective allele and probed them with a monoclonal antibody raised against the first 200aa of murine RECQL4 by Western blot (15). We found a truncated protein product of the predicted size for the R347X mutant in thymocyte extracts, though at a much lower intensity than the WT band (Fig 1D). In contrast, the G522Efs mutant protein could not be detected (Fig 1D), and even when ectopically overexpressed as a cDNA with an N-terminal 3xFlag tag, its expression was significantly lower than the R347X (Fig 1E).

To assess whether the truncated proteins had altered cellular localization, we generated N-terminal mCherry-tagged mouse RECQL4 fusion constructs. These were retrovirally infected into the murine osteoblastic Kusa4b10 cell line. Protein localization was analyzed by fluorescence microscopy. The full-length wild-type murine RECQL4 protein (WT) was predominantly localized in the nucleus as expected, with an apparent enrichment in the nucleolus. The ATP-dependent helicase inactive p.Lys525Ala (K525A) mutation, which we recently reported (15), had a similar localization to WT RECQL4 (Fig 1F). In contrast, the R347X protein, while also localized to the nucleus, was poorly incorporated in the nucleoli (Fig 1F). Interestingly, consistent with the weak protein expression *in vivo* (Fig 1D), the G522Efs protein was poorly expressed and difficult to detect (Fig 1F).

We further assessed whether the cellular localization patterns were shared with similar human RTS mutations. For this purpose, we utilized a human N-terminal EGFP-tagged WT, an ATP-helicase inactive K508A mutant, and created the C525Afs and R350Gfs mutations. These constitute the human homologues of, or map closely to, the murine mutations K525A, G522Efs, and R347X, respectively. We found comparable localization results between the human and murine proteins (Fig 1G).

Finally, it has been suggested that RECQL4 could localize to the mitochondria (23, 24). Using the fluorescent fusion proteins, we could not detect mouse or human RECQL4 (WT or mutant) in the cytoplasm nor overlapping with the mitochondria (S2 Fig A). To evaluate this result functionally, we assessed mitochondrial function using the Seahorse bioenergetic assay. To enable comparison of the different point mutations, we used HoxB8 immortalized myeloid progenitor cells (25) derived from *R26*-CreER *Recql4^fl/+^*, *R26*-CreER *Recql4^fl/K525A^, R26*-CreER *Recql4^fl/R347X^*, and *R26*-CreER *Recql4^fl/G522Efs^* and exposed them to tamoxifen for four days to delete the wild-type *Recql4* floxed allele. We found no difference in either basal or maximal oxygen consumption rate (OCR) between the non-tamoxifen and tamoxifen-treated groups (S2 Fig B-E), demonstrating that mutations in RECQL4 that significantly impact protein stability and function do not measurably affect mitochondrial respiration. Taken together, our results demonstrate that the murine RECQL4 mutants behave similarly to their human counterparts; and while the specific mutations impact their level of expression and subcellular localization differently, they do not measurably affect mitochondrial respiration.

### *Recql4^R347X/+^* and *Recql4^G522Efs/+^* heterozygosity leads to reduced bone mass phenotype

We previously reported that complete deletion of *Recql4* in the osteoblast lineage resulted in mice with shorter bones and reduced bone volume (22). RTS patients, however, present with compound heterozygous mutations that result in truncating proteins, rather than null alleles. Therefore, to assess the skeletal/bone phenotypes associated with RTS type 2 relevant RECQL4 mutations, we measured skeletal growth in the two viable heterozygous mutants. We could not evaluate homozygous mutants for either allele due to the developmental lethality previously described. The germline heterozygous animals are therefore most similar to the parents of RTS patients. For all genotypes, ten-week old male mice were analyzed. The heterozygous *Recql4^R347X/+^* mice were significantly smaller, while heterozygous *Recql4^G522Efs/+^* mice had a reduced body weight that did not reach statistical significance within the cohort assessed (p=0.076) (Fig 2A). We utilized Echo-MRI to investigate if the lower body weight was associated with a change in fat/lean mass proportion and found no differences, suggesting that the reduced mass was not likely to stem from changes in adiposity (S3 Fig). We then evaluated the tibial length by micro-computed tomography (micro-CT). Overall tibial length in both *Recql4^R347X/+^* and *Recql4^G522Efs/+^* mutants was not statistically different compared to either wild-type (littermate) controls or K525A homozygous males (Fig 2B). However, the mediolateral and anteroposterior widths measured by micro-CT in the midshaft tibia were significantly lower in both *Recql4^R347X/+^* and *Recql4^G522Efs/+^* mutants, indicating narrower tibiae in both genotypes (Fig 2C and 2D). For comparison, tibiae from the helicase-inactive K525A homozygous mice did not show differences in any of these parameters when compared to the WT controls.

**Fig. 2.**
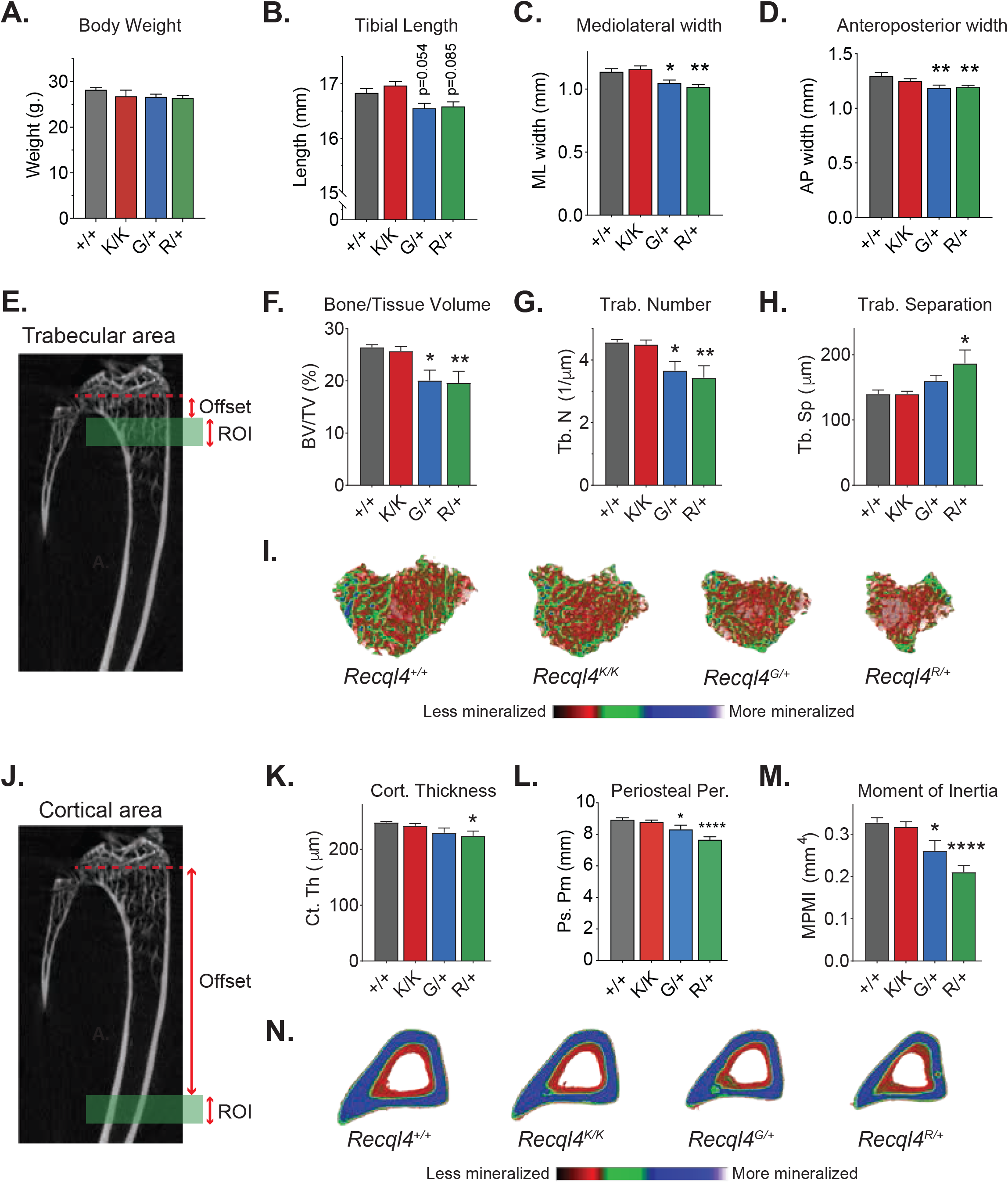
Germline truncating mutants G522Efs and R347X cause low bone mass and narrow bones. (A) Gross body weights of 10-week old males *Recql4^+/+^, Recql4^G522Efs/+^,* and *Recql4^R347X/+^* mice. Micro-CT measurements of (B) Tibial length, (C) Mediolateral width, and (D) Anteroposterior width from 10-week old males *Recql4^+/+^, Recql4^K525A/K525A^, Recql4^G522Efs/+^,* and *Recql4^R347X/+^* mice. (E) Trabecular region of interest beginning at 3.5% (Offset) distal to the growth plate and extending for 5% (ROI) of the total tibia length. (F) Trabecular bone volume. (G) Trabecular number. (H) Trabecular separation. (I) Representative images (Axial plane) of reconstructed trabecular bone with color-coded quantitative mineralization from germline *Recql4* mutants. (J) Cortical region of interest beginning at 40% (Offset) distal to the growth plate and extending for 5% (ROI) of the total tibia length. (K) Cortical thickness. (L) Periosteal perimeter. (M) Moment of inertia. (N) Representative images (Axial plane) of reconstructed cortical bone with color-coded quantitative mineralization from germline *Recql4* mutants. Data expressed as mean ± SEM, Ordinary one-way ANOVA. *P<0.05; **P<0.01; ***P<0.001; ****P<0.0001. +/+, n=7; K/K, n=10; G/+, n=6, R/+, n=7. Experiments were independently executed on separate cohorts, with results pooled for presentation. K=K525A; G=G522Efs; R=R347X.

We further looked at possible bone changes in trabecular microarchitecture and cortical morphology of WT and mutant 10-week old male mice. For trabecular analysis, we selected a region corresponding to the secondary spongiosa in the proximal metaphysis of the tibia (Fig 2E). The trabecular bone volume of *Recql4^R347X/+^* and *Recql4^G522Efs/+^* mice was significantly lower by 24% and 26%, respectively (Fig 2F). The trabecular number was also lower by 25% in the *Recql4^R347X/+^* and 20% in the *Recql4^G522Efs/+^* mice (Fig 2G). The trabecular separation was 25% greater in the *Recql4^R347X/+^* mice but not different in the *Recql4^G522Efs/+^* mice (Fig 2H). For cortical analysis, we assessed a region corresponding to the mid-diaphysis of the tibia (Fig 2J). Cortical thickness was 10% lower in the *Recql4^R347X/+^* mutants, whereas there was a slight (7%) but non-significant decrease in the *Recql4^G522Efs/+^* mice, compared to controls (Fig 2K). For both the *Recql4^R347X/+^* and *Recql4^G522Efs/+^,* the periosteal perimeter showed a reduction of 14% and 7%, respectively, when compared with littermate controls (Fig 2L), which was reflected in a lower moment of inertia for both groups (Fig 2M). This suggested a lower torsional rigidity and increased fracture risk in the germline heterozygous truncating mutant mice. In contrast, the non-truncating but ATP-binding helicase-inactive *Recql4^K525A/K525A^* mutants did not show any change in any trabecular or cortical parameter when compared to the WT control (Fig 2E-2H, 2J-2M). All morphological changes could be visualized in the color-coded 3D reconstructions (Fig 2I and 2N). Additional micro-CT parameters are provided in S4 Fig A-C. Collectively, these results demonstrate that heterozygous truncating mutations of RECQL4 in mice resulted in narrow bones and skeletal dysplasia.

Given that several studies have reported RTS patients with hematopoietic defects (26–28) and that there is an established reciprocal relationship between bone and hematopoiesis (29), we assessed whether the changes in skeletal parameters seen in the germline heterozygous mutant mice would impact hematopoiesis. Results showed a decrease in the cell hierarchy involved in myeloid development, which did not affect mature granulocytes or macrophages (S5 Fig). The remaining parameters assessed were normal. Therefore, a single copy of a truncating mutation in the presence of a retained WT allele is not sufficient to cause marked changes in hematopoiesis and is consistent with the reports from RTS patients and the apparent normality of hematopoiesis in their heterozygous parents. Taken together, these observations suggest that heterozygous truncating RECQL4 mutations cause a haploinsufficient low bone mass phenotype similar to that reported in RTS patients. Interestingly, the expression of a single copy of a full-length wild type RECQL4 is sufficient to maintain hematopoiesis, which indicates cellular differences between osteoblast lineage cells and blood-forming cells in sensitivity to *Recql4* gene dosage.

### Intrinsic defects in osteoblast lineage cells cause the low bone mass phenotype of RECQL4 truncating mutants

To determine whether the skeletal phenotypes seen in the germline heterozygous mutant mice were caused by defects in the osteoblast lineage, we crossed all our mutant models (*Recql4^K525A/+^, Recql4^R347X/+^* and *Recql4^G522Efs/+^*) to the *Osx*-Cre *Recql4^fl/fl^* mice (22, 30). This allowed us to delete the wild-type *Recql4* allele from the osteoblastic lineage, leaving only the mutant protein expressed. These osteoblast-restricted point mutant models were compared to *Osx*-Cre+ *Recql4*^fl/+^ mice to control for the known effects of the *Osx*-Cre transgene on bone homeostasis (31, 32) allowing comparison across all Cre+ models. Furthermore, this approach also bypassed the lethality of homozygous mutant mice and the assessment of cells only expressing truncated proteins in adult mice.

The analysis showed that only the *Osx*-Cre *Recql4^Δ/R347X^* mice had a lower body weight and tibial length compared with *Osx*-Cre *Recql4^Δ/+^* littermates (Fig 3A and 3B). The *Osx*-Cre *Recql4^Δ/G522Efs^* mice had a reduction in body weight, but it did not reach statistical significance (p=0.09) (Fig 3A). Additionally, trabecular analysis showed lower trabecular bone volumes by 13% for *Osx*-Cre *Recql4^Δ/G522Efs^* and 25% for *Osx*-Cre *Recql4^Δ/R347X^* (Fig 3C). Correspondingly, there was a 12% and 23% reduction in trabecular number, respectively, compared to controls (Fig 3D). Trabecular separation was 17% greater in the *Osx*-Cre *Recql4^Δ/R347^* mice only (Fig 3E). For cortical bone parameters, only mice carrying the R347X mutation showed an 8% lower periosteal perimeter and 25% lower mean polar moment of inertia consistent with an 8% reduction in mediolateral width and no change in anteroposterior width when compared to controls (Fig 3G-3K). Again, the K525A helicase-inactive mice did not show any detectable phenotypes when compared to controls. All morphological changes are illustrated in the 3D reconstructed images (Fig 3F and 3L). Additional micro-CT parameters are provided in S4 Fig D-F. In summary, these data demonstrate that truncating mutations of RECQL4 disrupt bone microstructure through defects intrinsic to the osteoblast lineage with the more severe phenotype seen in mice expressing the shortest truncated protein (R347X).

**Fig. 3.**
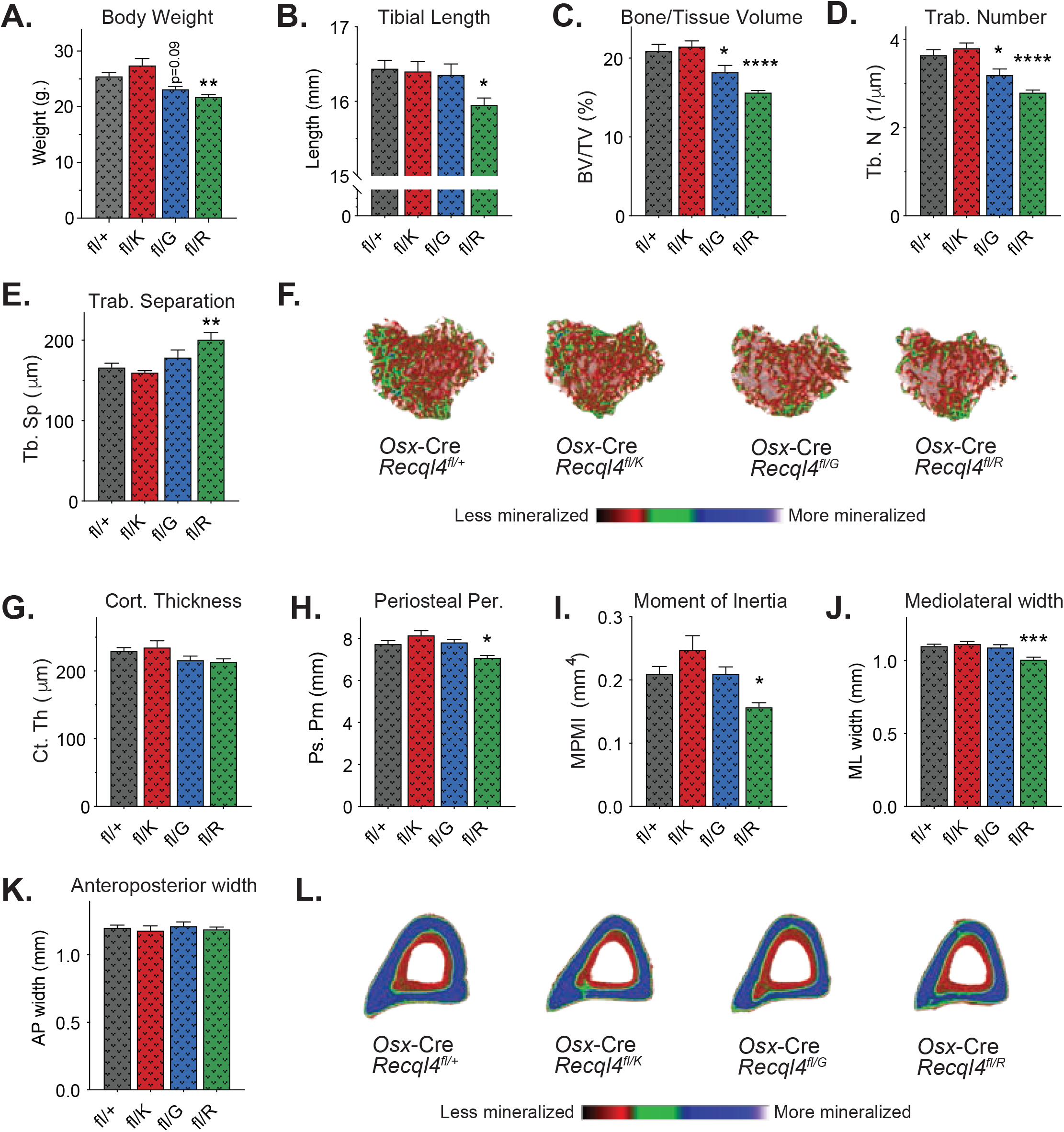
Expression of only the truncating mutations in pre-osteoblasts results in low bone mass. (A) Gross body weights of 10-week old males *Osx*-Cre *Recql4^fl/+^*, *Osx*-Cre *Recql4^fl/K525A^, Osx*-Cre *Recql4^fl/G522Efs^,* and *Osx*-Cre *Recql4^fl/R347X^* mice. (B) Tibial length. (C) Trabecular bone volume. (D) Trabecular number. (E) Trabecular separation. (F) Representative images (Axial plane) of reconstructed trabecular bone with color-coded quantitative mineralization from *Osx*-Cre *Recql4* mutants. (G) Cortical thickness. (H) Periosteal perimeter (I) Moment of inertia. (J) Mediolateral width. (K) Anteroposterior width. (L) Representative images (Axial plane) of reconstructed cortical bone with color-coded quantitative mineralization from *Osx*-Cre *Recql4* mutants. Data expressed as mean ± SEM, Ordinary one-way ANOVA. *P<0.05; **P<0.01; ***P<0.001; ****P<0.0001. fl/+, n=9; fl/K, n=6; fl/G, n=7, fl/R, n=7. Experiments were independently executed on separate cohorts, with results pooled for presentation. fl=Floxed; K=K525A; G=G522Efs; R=R347X.

### Compound heterozygous *Recql4* mutants tolerate ionizing radiation and do not develop osteosarcoma

Based on previous studies that show that RTS patients have compound heterozygous mutations with one allele more severely truncated than the other (4, 5), we generated compound heterozygous mouse lines combining the G522Efs or R347X mutations with the K525A. Although this approach does not fully mimic RTS, it allowed us to determine the *in vivo* effects of a truncated allele and a helicase-inactive allele. Surprisingly, the compound heterozygous *Recql4^G522Efs/K525A^* and *Recql4^R347X/K525A^* mice were viable, and pups were born at the expected Mendelian ratio (Fig 4A). Moreover, monitoring of aged cohorts demonstrated that *Recql4^G522Efs/K525A^* and *Recql4^R347X/K525A^* mice had a normal lifespan compared to WT mice without increased tumor incidence (Fig 4B). All genotypes developed a small number of spontaneous tumors affecting the liver, spleen, thymus, and intestine, however no osteosarcoma was detected in compound heterozygous mice (S1 Table).

**Fig. 4.**
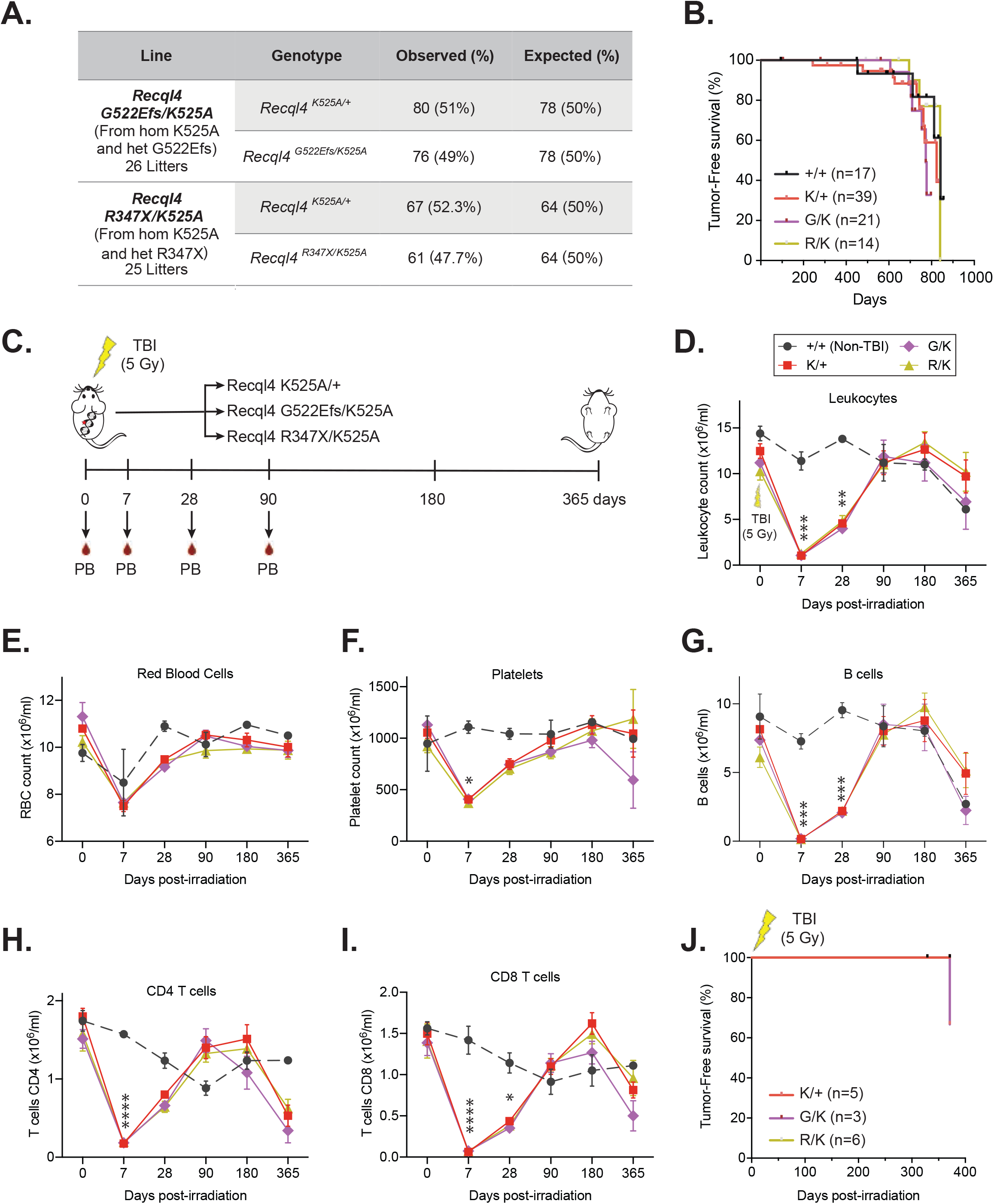
Compound heterozygous *Recql4* mutants tolerate a sublethal dose of ionizing radiation and do not develop osteosarcoma. (A) Breeding data from 26 litters of *Recql4^G522/K525A^* and 25 litters of *Recql4^R347X/K525A^* intercrosses. Observed and expected mendelian rates of the indicated genotypes are shown. No statistical significance was achieved. (B) Kaplan-Meier tumor-free survival plots of the indicated genotypes. +/+, n=17; K/+, n=39; G/K, n=21; R/K, n=14. (C) Schematic representation of experimental setup. Compound heterozygous mutants received a single dose of 5-Gy gamma-irradiation, and peripheral blood was assessed, and the animals monitored for tumor formation. Mice were euthanized at the last timepoint. Peripheral blood cell counts following irradiation: (D) Leukocytes; (E) Red blood cells; (F) Platelets; (G) B cells; (H) CD4 T cells; (I) CD8 T cells. (J) Kaplan-Meier tumor-free survival plots of the irradiated mice. K/+, n=5; G/K, n=3; R/K, n=6. Data expressed as mean ± SEM, Ordinary one-way ANOVA. *P<0.05; **P<0.01; ***P<0.001; ****P<0.0001. TBI=Total body irradiation; K=K525A; G=G522Efs; R=R347X.

Previous studies have reported increased sensitivity of RECQL4 mutant cell lines to ionizing radiation (33, 34). To assess the *in vivo* response of compound heterozygous mutant mice and whether this increased susceptibility to cancer formation, we treated a small cohort of mice with whole-body ionizing radiation. A single sub-lethal dose of 5Gy γ-irradiation was administered to 9-week old *Recql4^G522Efs/K525A^, Recql4^R347X/K525A^* and *Recql4^K525A/+^* mice as controls. We monitored the cohorts for one year, assessing peripheral blood parameters at several time points to evaluate hematologic recovery (Fig 5C). The efficacy of the radiation was demonstrated by a similar transient reduction in blood cell populations across all genotypes (Fig 4D-4I). After twelve months, all mice were euthanized and autopsies performed. We found that ionizing radiation did not result in abnormal/delayed hematopoietic recovery or failure nor did it result in tumor development with only one intestinal tumor found in a *Recql4^G522Efs/K525A^* mouse (Fig 4J). Collectively, these results demonstrate that a full-length, even helicase inactive RECQL4, is sufficient to rescue the lethality caused by truncating mutations. They further demonstrate that these compound heterozygous *Recql4* mutations are not sufficient to sensitize murine models to ionizing radiation or to accelerate cancer initiation.

**Fig. 5.**
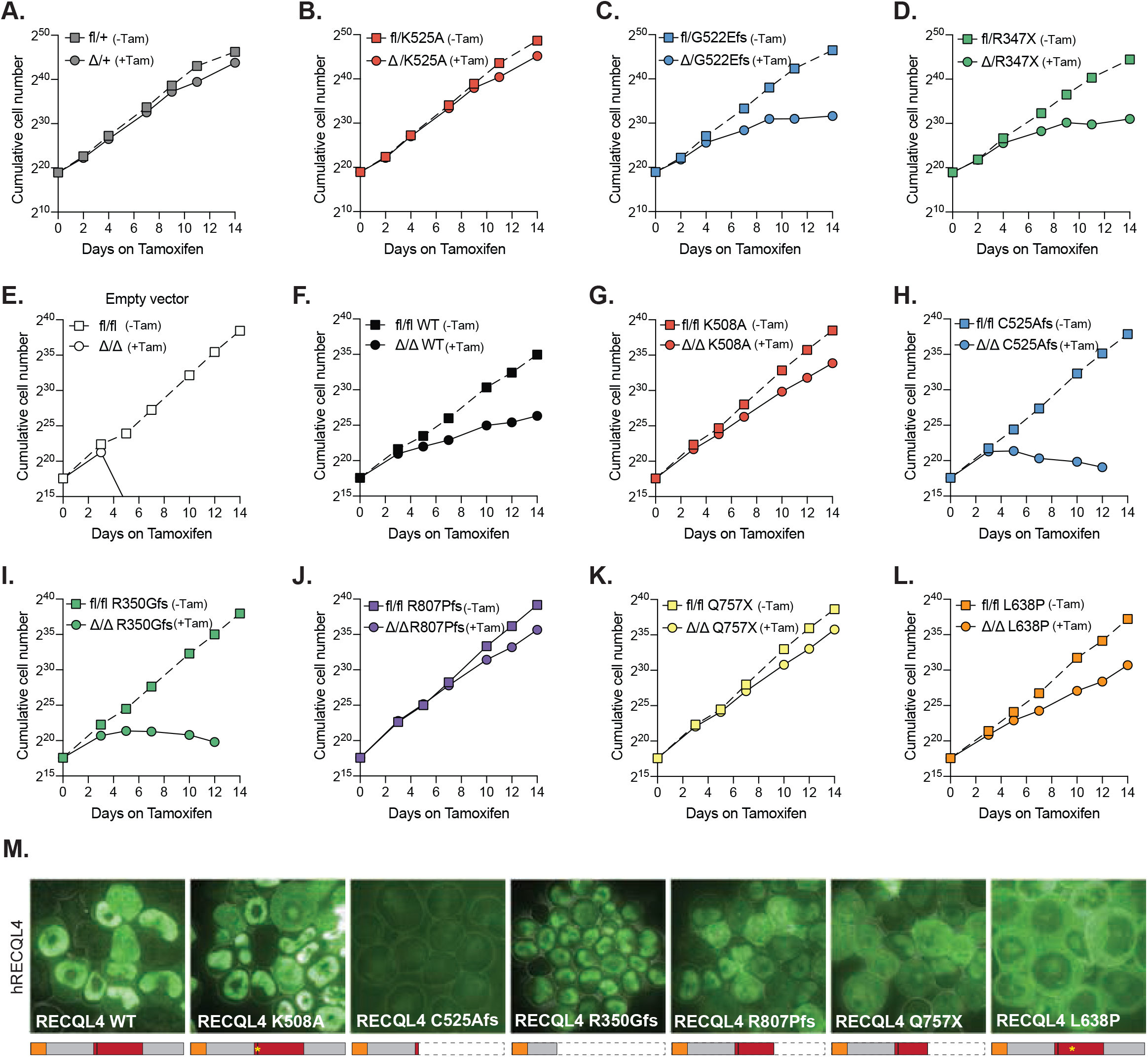
Truncating human RECQL4 mutations C525Afs and R350Gfs fail to rescue *Recql4* deletion and impede proliferation, similar to their murine homologs, while mutations that conserve the protein are better tolerated. Proliferation curves of HoxB8 immortalized *R26*-CreER^T2^ myeloid cells without (fl) and with (Δ) tamoxifen-mediated *Recql4* deletion in the following cell lines: (A) fl/+; (B) fl/K525A; (C) fl/G522Efs; (D) fl/R347X. Proliferation curves of HoxB8 immortalized *R26*-CreER^T2^ *Recql4^fl/fl^* control myeloid cells (E) in the presence or absence of tamoxifen, and EGFP hRECQL4 over-expressing cells: (F) Wild type; (G) K508A; (H) C525Afs; (I) R350Gfs; (J) R807Pfs; (K) Q757X; and (L) L638P. Dotted lines represent individual controls not treated with tamoxifen. (M) Microscopy of EGFP-hRECQL4 expression in HoxB8 cells with WT, K508A, C525Afs, R350Gfs, R807Pfs, Q747X, L638P. A schematic illustration of the expected protein products is outlined below each figure. Orange box represents the Sld2-like region, red box the helicase region. Cell proliferation assays using murine mutations were repeated two times using independent cell lines. Retroviral complementation assays with human constructs were performed two times using the same parental cell line.

### Human mutations causing severe truncations fail to rescue *Recql4* deletion recapitulating their murine homologs, while mutations that conserve the protein are better tolerated

To analyze the effects of different human RECQL4 mutations and how they compare to our murine models, we utilized the Hoxb8 immortalized myeloid progenitor cells. This is a relevant cell type, given the requirement of RECQL4 for maintenance of this cell population *in vivo* (35). First, we used this system to determine the effects of truncating murine *Recql4* mutations *in vitro*. Hoxb8 immortalized myeloid progenitor cell lines from *R26*-CreER^T2^ *Recql4^fl/+^*, *R26*-CreER^T2^ *Recql4^fl/K525A^, R26*-CreER^T2^ *Recql4^fl/R347X^*, and *R26*-CreER^T2^ *Recql4^fl/G522Efs^* mice were treated with 400nM/mL of 4-hydroxy-tamoxifen to induce Cre recombinase activity and deletion of the floxed wild type *Recql4* allele, leaving only the mutant allele expressed. We achieved successful deletion by day four, as confirmed by PCR for genomic recombination (S5 Fig). The presence of a single *Recql4* wild type allele (fl/+ cells treated with 4-hydroxy-tamoxifen to become Δ/+) did not interfere with the proliferation rates (Fig 5A), nor did the expression of a single helicase-inactive K525A allele (Fig 5B). However, when the floxed allele was deleted in the cells carrying the truncating mutations G522Efs and R347X, a marked decrease of cell proliferation was observed, consistent with the previously described *in vivo* phenotype (15) and demonstrating an essential requirement of the helicase and C-terminal domains deleted in these mutations (Fig 5C and 5D).

Next, we used this cell line model to determine the capacity of different human *RECQL4* mutations to rescue HoxB8 *R26*-CreER *Recql4^Δ/Δ^* myeloid cells, where both alleles of the endogenous murine *Recql4* were deleted. As expected, control fl/fl cells started dying at day four post-tamoxifen as they became *Recql4* null (Fig 5E and S6 Fig). Interestingly, when we expressed the wild type human RECQL4 protein, cell proliferation was not wholly restored, suggesting that overexpression of wild-type RECQL4 can be detrimental (Fig 5F). We further tested the ability of frequent human RTS associated *RECQL4* mutations, including the human equivalents to our murine germline mutations, to rescue Δ/Δ cell proliferation and viability. To do this, we used the same EGFP-fused human RECQL4 mutations: K508A, C525Afs, and R350Gfs (Fig 5G and 5H), described previously and engineered the p.Arg807ProfsTer38 (R807Pfs), p.Gln757* (Q757X) and p.Leu638Pro (L638P) mutations based on recurrent mutations described in RTS patients (4). The analysis showed that the helicase-inactive K508A mutant successfully rescued the Δ/Δ cells (Fig 5G), while both the C525Afs and R350Gfs human mutants did not (Fig 5H and 5I), similar to endogenous murine G522Efs and R347X (Fig 5C and 5D). Both R807Pfs and Q757X mutations rescued the *Recql4* deletion (Fig 5J and 5K), whereas the L638P achieved only partial rescue (Fig 5L).

Finally, we visualized the EGFP-RECQL4 localization in HoxB8 *R26*-CreER *Recql4^fl/fl^*. Similar to what we saw in Kusa4b10 cells (Fig 1G), the wild type human RECQL4, the K508A and the R350Gfs proteins were localized in the nucleus, while the C525Afs was poorly expressed (Fig 5M). On the other hand, the R807Pfs and Q757Pfs mutants displayed both nuclear and cytoplasmic localization, while the L638P mutant was located primarily in the cytoplasm. The sizes and expression levels of the predicted protein products were confirmed by Western blotting (S7 Fig). Collectively, these data demonstrate that not all mutations of RECQL4 are functionally equivalent. Mutations that result in severely truncated protein products, due to early stop codons or frameshifts, are more likely to affect transcript stability and localization and are detrimental to both cellular and organismal health. In contrast, mutations that conserve most of the protein are largely tolerable.

## Discussion

Since Kitao *et al.* first cloned the *RECQL4* gene more than twenty years ago (36), some inroads have been made in the understanding of these mutations and their contribution to RTS. It is now known that the majority of mutations reported in RTS patients are compound heterozygous mutations, containing at least one truncated allele and mainly impacting the helicase domain (4, 5). This mutational spectrum implied that defects in the helicase region might be the main reason for the phenotypes of RTS. However, we recently showed that mice with a homozygous knock-in mutation that specifically inactivates RECQL4 ATP-dependent helicase activity were strikingly normal in terms of embryonic development, hematopoiesis, and DNA damage repair (15). In the same study, by using a conditional deletion model that allowed the assessment of the effects of mutations after deleting the wild type *Recql4* allele, we found that mice carrying truncating, but not helicase-inactive, mutations developed bone marrow failure (15). This confirmed the deleterious effects of truncating mutations, which led us to investigate the impact of these mutations in other systems.

Assessing the reported *RECQL4* mutations (1, 4, 5), RTS patients presenting with two severe truncating mutations are extremely rare. Here, we generated two *Recql4* mutant models with truncating mutations that closely map to those reported in RTS patients, affecting the helicase and C-terminal domain. We did not recover viable homozygous germline mutant pups from either allele. These results are consistent with the human data and suggest that having two severely truncating mutations of *RECQL4* is not tolerated and that there is an essential developmental role for the deleted domains. Furthermore, in addition to lacking essential domains, there is the potential for these mutations to yield aberrantly expressed and localized protein, further contributing to the phenotypes we observed. When we assessed protein levels of the truncating mutants, we were not able to detect the predicted truncating protein product from the G522Efs mutation. This could be either the consequence of nonsense-mediated mRNA decay or proteasomal degradation. The C525Afs human mutation, which maps closely to the murine G522Efs mutation, showed similar results. On the other hand, the R347X mutation produced a short but stable protein that localized to the nucleus, albeit with reduced nucleolar intensity compared to WT or K525A protein. Interestingly, this truncated protein retained the nuclear targeting sequence 1 (NTS1, 37-67aa) (37) but lacked the other of the two predicted nucleolar localization signals. The R350Gfs human mutation (mapping closely to the R347X) had similar localization and expression. Taken together, we have shown that *Recql4* truncating mutations affect protein stability and subcellular localization differently and that this phenotype is reproduced using comparable human mutations.

The skeletal system is severely impacted in the majority of RTS patients. A study of 28 RTS subjects examined by radiologic survey found that up to 75% had some form of skeletal abnormality, including abnormal metaphyseal trabeculation, brachymesophalangy, thumb or radial agenesis or hypoplasia (3). Several additional studies have associated loss of RECQL4 with a more systemic skeletal involvement with a high proportion of patients reporting low bone density (3, 38-41). However, to our knowledge, no studies have mapped skeletal defects to specific RECQL4 mutations. We found that heterozygous expression of the truncating alleles was sufficient to cause low trabecular bone mass, impaired growth of cortical bone, and narrower bones, compared to WT controls. Furthermore, we found that truncating mutations affect normal skeletal formation by causing defects in the osteoblast lineage. These findings were comparable to findings we previously reported in an osteoblast lineage restricted knock-out, which showed that complete deletion of *Recql4* in the osteoblast lineage led to reduced bone volume and defects in osteoblast proliferation and maturation (22). This suggested that at least for bone development, having a truncated RECQL4 protein is equivalent to having no RECQL4 at all, which highlights the critical function of the deleted domains in bone homeostasis. Interestingly, an osteochondral-lineage-specific mouse model reported more severe findings than ours. Cao *et al.* used *Prx1*-Cre to delete *Recql4* in early mesenchymal progenitor cells of the limb buds and described a 50% reduction in bone volume and cortical bone area (40). While RECQL4 is clearly needed in the earlier skeletal cell populations, it is important to consider that our models use truncating heterozygous mutations, instead of null alleles. Thereby, any remaining RECQL4 protein in the pre-osteoblast population of our mutants, might be sufficient for partial function at this stage and explain the phenotypic differences between models. Lastly, we observed that the R347X mutation, which produced a stable yet the shortest predicted protein, led to a more severe bone phenotype than the poorly expressed G522Efs mutation. This suggested that the severity of the defects was proportional to the severity of the truncation, irrespective of protein expression. Furthermore, the fact that the largest truncation caused the most severe bone phenotype suggests that RECQL4 gene dosage is a critical regulator of bone mass, something relatively unknown until now. Collectively, these results demonstrate that heterozygous truncating mutations of RECQL4 cause a haploinsufficient low bone mass phenotype through defects in the osteoblast lineage. Although RTS patients generally present with compound heterozygous mutations, these results highlight the importance of having a full-length protein for bone development. Furthermore, they raise concerns regarding the osteoporosis status of the parents of RTS patients, which warrants further investigation.

A characteristic feature of syndromes associated with mutations in RECQL4 is cancer predisposition, particularly osteosarcoma, cutaneous epithelial tumors, and hematological malignancies (2, 42, 43). A recent study analyzed pediatric patients with cancer and identified a significant enrichment in heterozygous *RECQL4* loss-of-function variants in those who presented with osteosarcoma (44). This raised the question whether the presence of compound heterozygous *Recql4* mutations in mice is sufficient to initiate tumorigenesis. We found no differences in tumor burden or spectrum in our compound heterozygous mutants compared to wild type controls, even after exposure to γ-irradiation. Furthermore, although previous studies have reported ionizing radiation as a risk factor for the development of sarcomas (45, 46), and truncated RECQL4 products have been associated with hypersensitivity to this agent (47), we did not observe either hematopoietic failure nor osteosarcoma development in our irradiated cohort. It is possible that the non-truncating but helicase-inactive allele is sufficient to rescue these phenotypes and that truncating mutations in both alleles are necessary for tumor initiation. However, this could not be addressed in this study given the developmental lethality seen in our homozygous truncating mice. Another possibility is that additional genes are involved in the development of these RTS phenotypes.

Lastly, we established a tractable *in vitro* cell line model, which allowed us to examine the cellular consequences of different *RECQL4* mutations. We found that cells carrying the murine truncating G522Efs and R347X mutations developed a proliferation defect after deletion of the floxed wild-type *Recql4* allele. Similarly, the closely related human C525Afs and R350Gfs mutations failed to rescue the lethality caused by *Recql4 deletion*. On the other hand, the murine ATP-dependent helicase inactive mutation (K525A) did not demonstrate a proliferation defect, and its human counterpart (K508A) successfully rescued *Recql4* deletion. These results confirm that truncating, but not helicase-inactive mutations are pathogenic and that our murine mutations are useful surrogates for understanding the functions of human disease-associated *RECQL4* mutations. When using this system to analyze other human mutations, we found that cells overexpressing the human wild type protein could not completely rescue *Recql4* deletion, suggesting that overexpression of RECQL4 is not well tolerated. In fact, several studies have correlated overexpression of *RECQL4* with the development of malignancies (48–53). We also found that the L638P mutation, which resulted in a stable full-length product with cytoplasmic localization, could not fully rescue the *Recql4* deletion. This demonstrates that in order for RECQL4 to function effectively, it needs to be located within the nucleus. Finally, the R807Pfs and Q757X mutations, which have been found in RTS patients with osteosarcoma and lymphomas (4), were able to rescue the *Recql4* deletion. This indicates that small C-terminal deletions do not severely affect viability, unlike larger deletions that include both the helicase and the C-terminal domains. However, their association with malignancies remains unknown. By comparing different mutations in the same genetic context, this system allowed us to conclude that the different mutations have distinct consequences for RECQL4. While some mutations create unstable proteins, some alter its localization without grossly affecting protein stability. Overall, the level of the truncation appeared to show the strongest correlation with the severity of the phenotype. Given the increasing number of somatic *RECQL4* mutations reported in sporadic cancers, this *in vitro* system can serve as a platform to assess the impact of *RECQL4* mutations at a cellular level.

In conclusion, truncating RECQL4 mutations affect protein stability and localization, contributing to the development of an osteoporosis-like phenotype through defects in early osteoblast progenitors in mice. However, they are not sufficient to promote tumorigenesis, even after exposure to irradiation. Future studies should focus on the identification of genes that co-operate with RECQL4 in normal development and tumorigeneses. These will allow a better understanding of the genetic landscape of RTS and permit the generation of more comprehensive models.

## Materials and Methods

### Ethics Statement

All animal experiments conducted for this study were approved by the Animal Ethics Committee of St. Vincent’s Hospital, Melbourne, Australia (#007/14, 011/15, and 015/17). Animals were euthanized by cervical dislocation or CO_2_ asphyxiation.

### Mice

The chemically (ENU, N-ethyl-N-nitrosourea) induced *Recql4^R347X^* point mutation was provided by the Australian Phenomics Facility (APF, Canberra, Australia; allele IGL01809). *Recql4^G522Efs^* mice were identified during CRISPR/Cas9 targeting to generate the previously described *Recql4^K525A^* mutation (15) by the Mouse Engineering at Garvan/ABR (MEGA) service (Garvan Institute, Darlinghurst, Australia). This allele is on a C57BL/6 background and carried a 2bp insertion (GA) after the T521 codon. The *Osx*-Cre, *Rosa26*-eYFP, and *Recql4^K525A^* mutant animals have been previously described (15, 22, 30). *Rosa26*-CreER^T2^ mice were originally purchased from The Jackson Laboratory (B6.129-*Gt(ROSA)26Sor^tm1(cre/ERT2)Tyj^*/J, Stock Number: 008463) and have been previously described (35). The ENU derived mutant (*Recql4^R347X^*) was backcrossed to C57BL/6 at least six times and evaluated across multiple generations. All lines were on a C57BL/6 background. All animals were housed at the BioResources Centre (BRC) at St. Vincent’s Hospital. Mice were maintained and bred under specific pathogen-free conditions with food and water provided *ad libitum.*

All mouse lines are available from the Australian Phenome Bank (APB; https://pb.apf.edu.au/). Strain identification numbers/names are: R347X (APB ID#7986); *R26*-CreER *Recql4^fl/fl^* (APB ID#7263); *Osx*-Cre *R26*-eYFP *Recql4^fl/fl^* (APB ID#7886); K525A (strain name: C57BL/6-Recql4<tm4Crw>) and G522Efs (strain name: C57BL/6-Recql4<tm5Crw>).

### Cloning of mCherry and EGFP RECQL4 proteins and Retroviral production

Human N terminal EGFP fused RECQL4 and EGFP fused RECQL4^K508A^ (provided by T. Enomoto, Musashino University, Tokyo, Japan; (54)) were cloned into MSCV-puro (35). Human mutations R807Pfs, Q757X, L638P, C525Afs, and R350Gfs, were created by gBlock (IDT) replacement of a wild-type fragment of EGFP-RECQL4 in the plasmid MSCV-puro with a mutant fragment. Murine mCherry fused RECQL4 was assembled from a codon-optimized synthetic m*Recql4* cDNA (GeneArt, Life Technologies), placed in frame with an N-terminal mCherry cDNA (gBlock, IDT) in MSCV-puro. Mouse mutations were created by gBlock (IDT) replacement of the required fragment of *Recql4*. All constructs contain full-length cDNAs, including those coding for truncating mutations. All mutations were confirmed by Sanger sequencing. Retrovirus was produced by transient transfection of 293T cells using calcium phosphate mediated transfection with an ecotropic packaging plasmid (35).

### Genotyping

Genotyping of the G522Efs mutants was determined by PCR using the following primers: mRecql4 K525A MO36-R3: 5’-AGAACATTGGGCATTCGGC-3’ and mRecql4 K525A MO36-F9: 5’-TAGACCTTATGAAACCTCAAAGCC-3’ to obtain a 591bp product, which was then digested with *MslI* (NEB) to generate three fragments of 347, 175 and 71bp for the G522Efs mutant; or two fragments of 416 and 175bp for the WT. Genotyping of the K525A mutants has been previously described and used the same primers and restriction enzyme as the G522Efs mutation with the difference that this resulted in three fragments of 361, 175, and 55bp (15). The presence of the R347X mutation was determined by KASP (competitive allele-specific PCR) technology (LGC) with facility provided primers: 5’-GAAGGTGACCAAGTTCATGCTAAAGCGTTTGTTTTTCATGTTGAGTCG-3’, 5’-GAAGGTCGGAGTCAACGGATTCAAAGCGTTTGTTTTTCATGTTGAGTCA-3’, and reverse primer 5’-GCTTCCCTAGACAGAGGGAACTATA-3’ used according to manufacturer instructions.

### Protein extraction and Western blotting

Thymocyte lysates from germline *Recql4^R347X/+^* and *Recql4^G522Efs/+^* were prepared in RIPA buffer (50mM Tris, 150mM NaCl, 1% NP-40, 0.5% sodium deoxycholate, 0.1% SDS, pH8.0) plus Complete protease inhibitor (Roche) and PhosStop (Roche) tablets. Protein was quantified using the Pierce BCA protein assay kit (Thermo Fisher Scientific) on an Enspire multimode plate reader (Perkin Elmer). Lysates from HoxB8 immortalized (25) *R26*-CreER^T2^ *Recql4^fl/G522Efs^, R26*-CreER^T2^ *Recql4^fl/R347X^* and *R26*-CreER^T2^ *Recql4^+/+^* transduced with MSCV puro 3xFlag-RECQL4 were prepared in sample buffer (2×10^6^ cells in 100μl). 50μg of whole protein extracts from thymocytes and 50μl from myeloid cells were loaded on pre-cast NuPAGE™ BOLT 8% Bis-Tris polyacrylamide gels (Invitrogen) and transferred onto Immobilon-P PVDF membranes (Merck Millipore). Membranes were blocked with 5% milk in Tris-buffered saline with tween (TBST) and incubated at 4°C overnight with rat monoclonal anti-mouse RECQL4 antibody (clone 3B10, detects mouse; or clone 3B1, detects both human and mouse) (15), mouse anti-Actin (Sigma Aldrich, A1978), or anti-Flag antibody (Sigma Aldrich). Membranes were then probed with HRP-conjugated goat anti-rat (Thermo Fisher Scientific, 31470) or anti-mouse (Thermo Fisher Scientific, 31444) secondary antibodies and visualized using ECL Prime Reagent for chemiluminescent detection on Hyperfilm ECL (Amersham). The predicted molecular weight for the truncated proteins is 62.4kDa for G522Efs, 65.6kDa for 3xFlag-G522Efs, 38.1kDa for R347X, and 41.3kDa for 3xFlag-R347X.

### Live cell imaging and image processing of RECQL4 fusion proteins

Transduced osteoblast-like Kusa4b10 cells were plated in 10-cm dishes, selected with puromycin, and grown to sub-confluency. Single plane images were acquired on an inverted fluorescent microscope (Olympus IX81) with a 40X objective (LUCPLFLN 40X) and were recorded with a Retiga-EXi 12 Bit CCD camera (QImaging). Image processing and analysis were done using MetaMorph (Molecular Devices) and Adobe Photoshop. HoxB8 immortalized myeloid cells were concentrated by centrifugation and 5μl of cell suspension dispensed on a slide before image acquisition with a 60X objective (UPLANAPO 60X water immersion). For mitochondrial staining, 250μL of Mitochondrial Staining Solution (CytoPainter MitoBlue Indicator Reagent (ab219940), 1:500 diluted in Hank’s Balanced Salt solution + 20 mM HEPES buffer (HHBS)) was added to Kusa4b10 cells grown on coverslips in 250μL DMEM in a 24-well plate and incubated for 30 minutes to 2 hours in a 37ºC/5% CO_2_ incubator. Coverslips were washed twice with HHBS, and live cells were imaged as described above.

### Seahorse XF24 Extracellular Flux analyzer

Hoxb8 immortalized (25) *R26*-CreER^T2^ *Recql^fll/+^* (control), *R26*-CreER^T2^ *Recql4^fl/K525A^, R26*-CreER^T2^ *Recql4^fl/G522Efs^,* and *R26*-CreER^T2^ *Recql4^fl/R347X^* myeloid cells were maintained in IMDM, 10% FBS (non-heat inactivated) and 1% GM-CSF containing media (BHK-HM5 cell-conditioned media). The cells were treated for four days with 400nM 4-hydroxy tamoxifen (Merck Millipore) then genotyped to confirm complete recombination. To adhere myeloid (suspension) cells to the XF24 cell culture plate, wells were first coated with 100μl of RetroNectin solution (32μg/ml in PBS, Takara Bio), incubated for 2 hours at room temperature, then blocked with 200μl of 2% BSA in PBS, and washed with PBS. Cells were then seeded at 120,000 cells/well and spun at 1100g for 20 seconds. Cell culture media was replaced with non-buffered DMEM base media (Seahorse bioscience) containing 25mM glucose, 1mM sodium pyruvate, glutamine at pH 7.4, and incubated for 1 hour at 37°C in a non-CO_2_ incubator. The oxygen consumption rate (OCR) was measured in a Seahorse XF24-3 Flux Analyzer. Cells were assayed with a 2-min mix/2-min wait/5-min measurement cycle for three baseline measurements followed by three cycles after each injection of four compounds affecting bioenergetics: 0.5μM oligomycin (Complex V inhibitor; Sigma, St. Louis, MO, USA), 0.7μM carbonyl cyanide 4-(trifluoromethoxy)phenylhydrazone (FCCP; ΔΨm uncoupler; Sigma), 3.6μM antimycin A (Complex III inhibitor; Sigma), and 6μM rotenone (Complex I inhibitor; Sigma). After completion of the analysis, Cyquant (Life technologies) was used to normalize measurements to cell number in the corresponding wells.

### Micro-Computed tomography (micro-CT) 3D analysis of tibia

Tibiae were collected from mutant mice and their littermate controls; the attached soft tissue was removed carefully, and tibiae were fixed in 2% paraformaldehyde overnight, which was then replaced by 70% ethanol. Tibia morphology and microarchitecture was analyzed by ex-vivo micro-CT on the left tibia wrapped in 70% ethanol-soaked gauze within a cryovial by using a Skyscan 1076 system (Bruker MicroCT, Kontich, Belgium). Images were acquired at 9μm pixel size, 0.5mm aluminum filter, 44kV voltage, 220μA current, 2300ms exposure time, 0.5° rotation, 1 frame averaging. Image slices were reconstructed by NRecon (Bruker, version 1.6.10.2) using the following settings: 36% beam-hardening correction, 6 ring artifact correction, 1 smoothing, and no defect pixel masking. The reconstructed images were analyzed with software programs Dataviewer (Bruker, version 1.4.4), CTan (Bruker, version 1.15.4.0), and CTVox (Bruker, version 2.4.0). The trabecular and cortical region of interest (ROI) was determined by identifying the start of the mineralized zone of the proximal growth plate and calculating a distance equal to 3.5% and 40% of the total tibial length, respectively. From that point, a further 5% of the total tibial length was analyzed as the secondary spongiosa trabecular ROI and a 5% as the cortical ROI. Analysis of bone structure was completed by adaptative thresholding in CTan, which was determined by performing automatic thresholding on 3 samples from each experimental group resulting in threshold values of 50-255 for trabecular bone and 90-255 for cortical bone. Representative images of reconstructed trabecular and cortical bone with color-coded quantitative mineralization were made of the specimen whose value was closest to the group mean using the trabecular bone volume and cortical thickness parameters.

### Peripheral blood analysis

Peripheral blood (approximately 100μl) was obtained via retro-orbital bleeding. 25μl of blood was mixed with 75μl of PBS to obtain cell counts on a hematological analyzer (Sysmex KX-21N, Roche Diagnostics). The remaining blood was red blood cell-depleted using hypotonic lysis buffer (150mM NH_4_Cl, 10mM KHCO_3_, 0.1mM Na_2_EDTA, pH7.3) and resuspended in 150μl of FACS buffer for flow cytometry analysis.

### Flow cytometry analysis

Bones were flushed, spleens and thymus crushed, and single-cell suspensions were prepared in FACS buffer. Antibodies against murine Ter119, CD71, B220, IgM, CD43, CD19, CD21, CD23, Mac-1, Gr1, F4/80, CD4, CD8, TCRβ, CD25, CD44, Sca-1, c-Kit, CD34, FLT3, FcγRII/III (CD16/32), CD41, CD105, CD150, either biotinylated or conjugated with phycoerythrin, phycoerythrin-Cy7, peridinin chlorophyll protein-Cy5.5, allophycocyanin, allophycocyanin eFluor780, eFluor660 or eFluor450 were all obtained from eBioscience, BioLegend or BD Pharmingen (S2 Table) (15, 35, 55, 56). Biotinylated antibodies were detected with streptavidin-conjugated with Brilliant Violet-605. 30,000-500,000 cells were acquired on a BD LSRIIFortessa and analyzed with FlowJo software Version 9 or 10.0 (Treestar).

### Cell proliferation assays

Hoxb8 immortalized (25) *R26*-CreER^T2^ *Recq4l^fl/+^*, *R26*-CreER^T2^ *Recql4^fl/G522Efs^,* and *R26*-CreER^T2^ *Recql4^fl/R347X^* cells were maintained in IMDM, 10% FBS (non-heat inactivated), 1% Pen/Strep, 1% L-Glutamine, and 1% GM-CSF containing media (BHK-HM5 cell-conditioned media). The cells were treated for 14 days with 400nM 4-hydroxy tamoxifen (Merck Millipore) then genotyped to confirm complete recombination. Cells were counted with Trypan blue using a Countess II automated counter (Thermo Fisher Scientific) and then split every two-three days.

### Retroviral transduction and complementation

Hoxb8 immortalized *R26*-CreER^T2^ *Recql4^fllfl^* cells were maintained in IMDM, 10% FBS (non-heat inactivated), 1% Pen/Strep, 1% L-Glutamine, and 1% GM-CSF containing media (BHK-HM5 cell-conditioned media). Exponentially growing Hoxb8 *R26*-CreER^T2^ *Recql4^fl/fl^* cells (100,000 cells/mL) were spin-infected with EGFP-RECQL4 retrovirus in a 1:1 ratio at 1100g for 90 minutes in 8μg/mL polybrene. Two days after infection, cells were selected with puromycin (0.25μg/mL) for four days and then expanded. Cells were then treated for 14 days with 400nM 4-hydroxy tamoxifen (Merck Millipore) and genotyped to confirm complete recombination. Cells were counted with Trypan blue using a Countess II automated counter (Thermo Fisher Scientific) and then split every two-three days.

### Statistical analysis

To determine statistical significance, Kaplan-Meier survival plots and ordinary one-way ANOVA tests were conducted in Prism software version 8 (GraphPad; San Diego, CA, USA). Throughout this study, significance is indicated using the following convention: *P<0.05; **P<0.01; ***P<0.001; ****P<0.0001, and data is presented as mean ± S.E.M. Furthermore, the number of samples used for each experiment is described in the corresponding figure legends

## Acknowledgments

We thank R. Brink and the Mouse Engineering Garvan/ABR (MEGA) Facility (Garvan Institute, Sydney, Australia) for the generation of the G522Efs allele; the Australian Phenomics Facility and the Australian National University (Canberra, Australia) for their technical expertise and provision of the R347X allele; S Galic and L Murray-Segal (St Vincent’s Institute) for training on Echo-MRI; T. Enomoto (Musashino University, Tokyo, Japan) for providing human EGFP-RECQL4 WT and K508A constructs; M. Kamps (University of California San Diego, USA) for providing the Hoxb8 vectors used to generate cell lines; D. Thorburn (Murdoch Children’s Research Institute and University of Melbourne, Australia) and J. Heierhorst (St Vincent’s Institute) for comments on the manuscript.

## Author contribution statement

**Conceptualization:** Wilson Castillo-Tandazo, Monique F. Smeets, Carl R. Walkley.

**Funding acquisition:** Carl R. Walkley.

**Investigation:** Wilson Castillo-Tandazo, Ann E. Frazier, Monique F. Smeets, Carl R. Walkley.

**Methodology:** Wilson Castillo-Tandazo, Ann E. Frazier, Natalie A Sims, Monique F. Smeets, Carl R. Walkley.

**Project administration:** Monique F. Smeets, Carl R. Walkley.

**Supervision:** Monique F. Smeets, Carl R. Walkley.

**Visualization:** Wilson Castillo-Tandazo, Monique F. Smeets, Carl R. Walkley.

**Writing – original draft:** Wilson Castillo-Tandazo, Monique F. Smeets, Carl R. Walkley.

**Writing – review & editing:** Wilson Castillo-Tandazo, Ann E. Frazier, Natalie A. Sims, Monique F. Smeets, Carl R. Walkley.

## Supporting Information

**S1 Fig.**
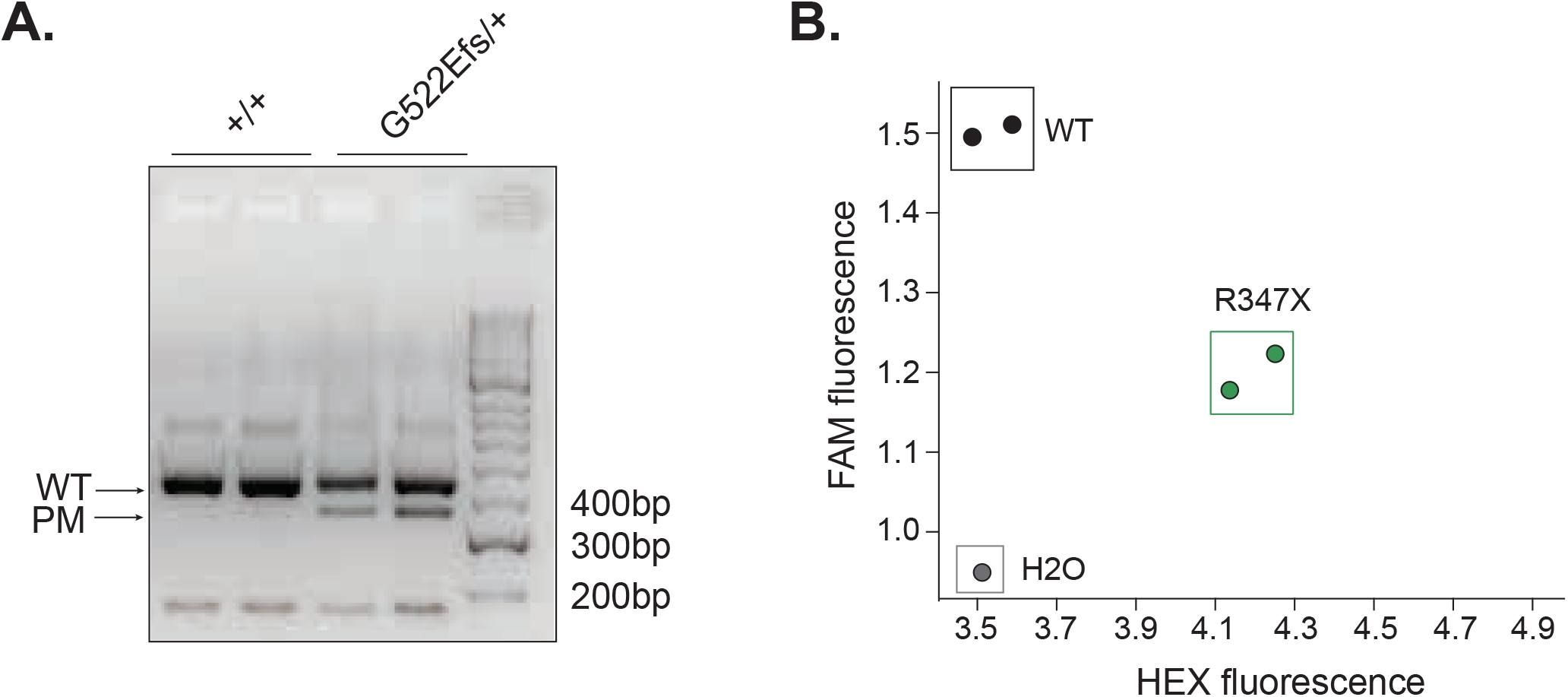
Genotyping of point mutant alleles. (A) Genomic DNA PCR plus Msl1 digest of *Recql4^+/+^* and *Recql4^G522Efs/+^.* Two independent mice per genotype. (B) KASP genotyping of *Recql4^+/+^* and *Recql4^R347X/+^.* HEX positive represents the R347X allele; FAM positive represents wild type allele. Two independent mice per genotype.

**S2 Fig.**
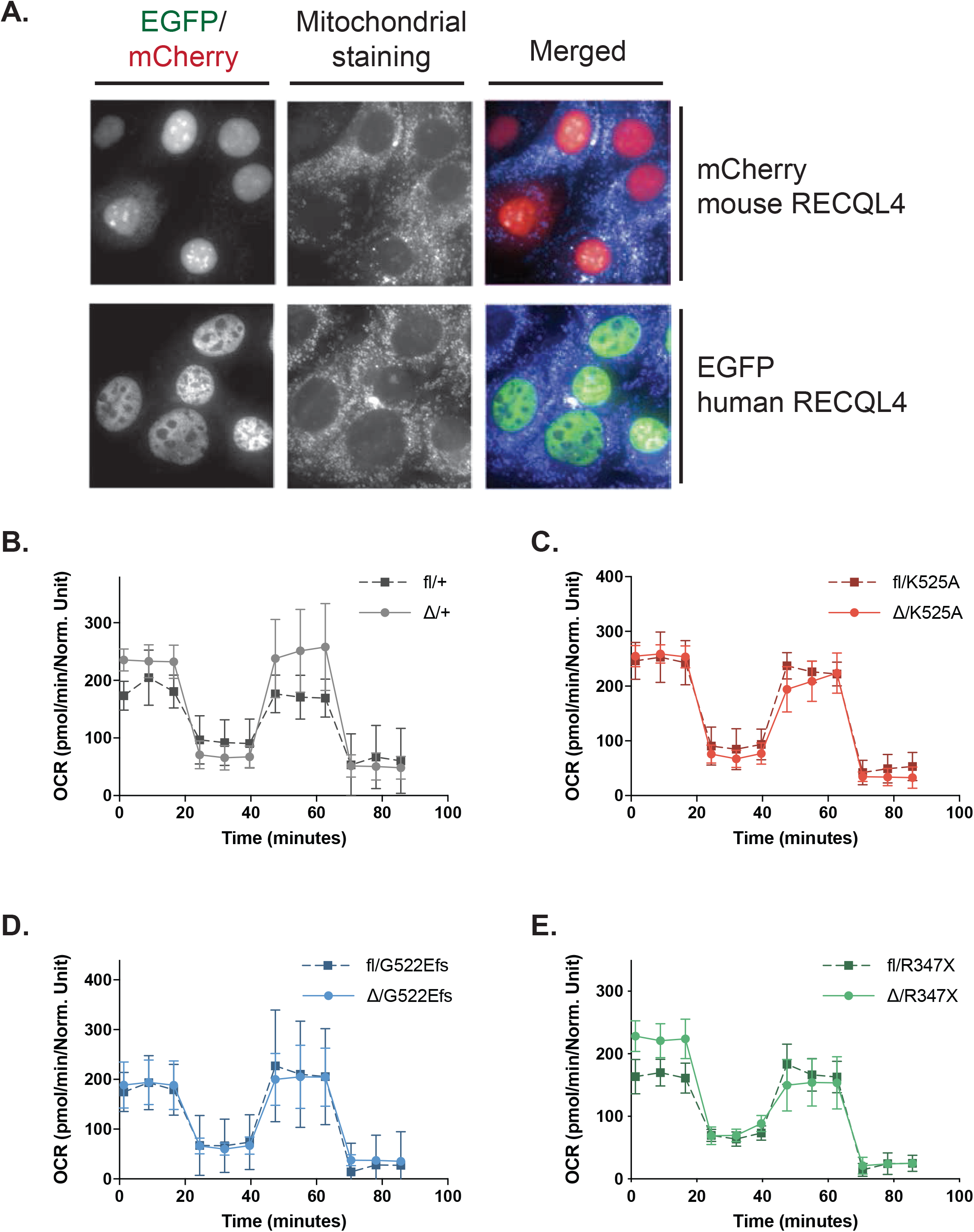
RECQL4 does not localize with mitochondria and truncating RECQL4 mutations do not affect mitochondrial respiration. (A) Localization of mCherry (red) fused mouse RECQL4 and EGFP (green) fused human RECQL4 in Kusa4b10 cells. In the same cell lines, mitochondrial localization was determined by staining with Mitochondrial Staining Solution (MitoBlue, 1:500 diluted in HHBS). Images were colorized and merged. (B-E) Seahorse XF24-3 instrument analysis of oxygen consumption rate (OCR) as a reflection of mitochondrial respiration in HoxB8 immortalized *R26*-CreER^T2^ myeloid cells after tamoxifen-mediated *Recql4* deletion (day 4 after tamoxifen addition) in the following cell lines: (B) Δ/+; (C) Δ/K525A; (D) Δ/G522Efs; (E) Δ/R347X. Dotted lines represent isogenic controls, not treated with tamoxifen. Data expressed as mean ± SEM. No statistical significance achieved between the tamoxifen-treated and non-treated groups. The experiment was performed using triplicate wells and was independently executed two times using the same cell lines.

**S3 Fig.**
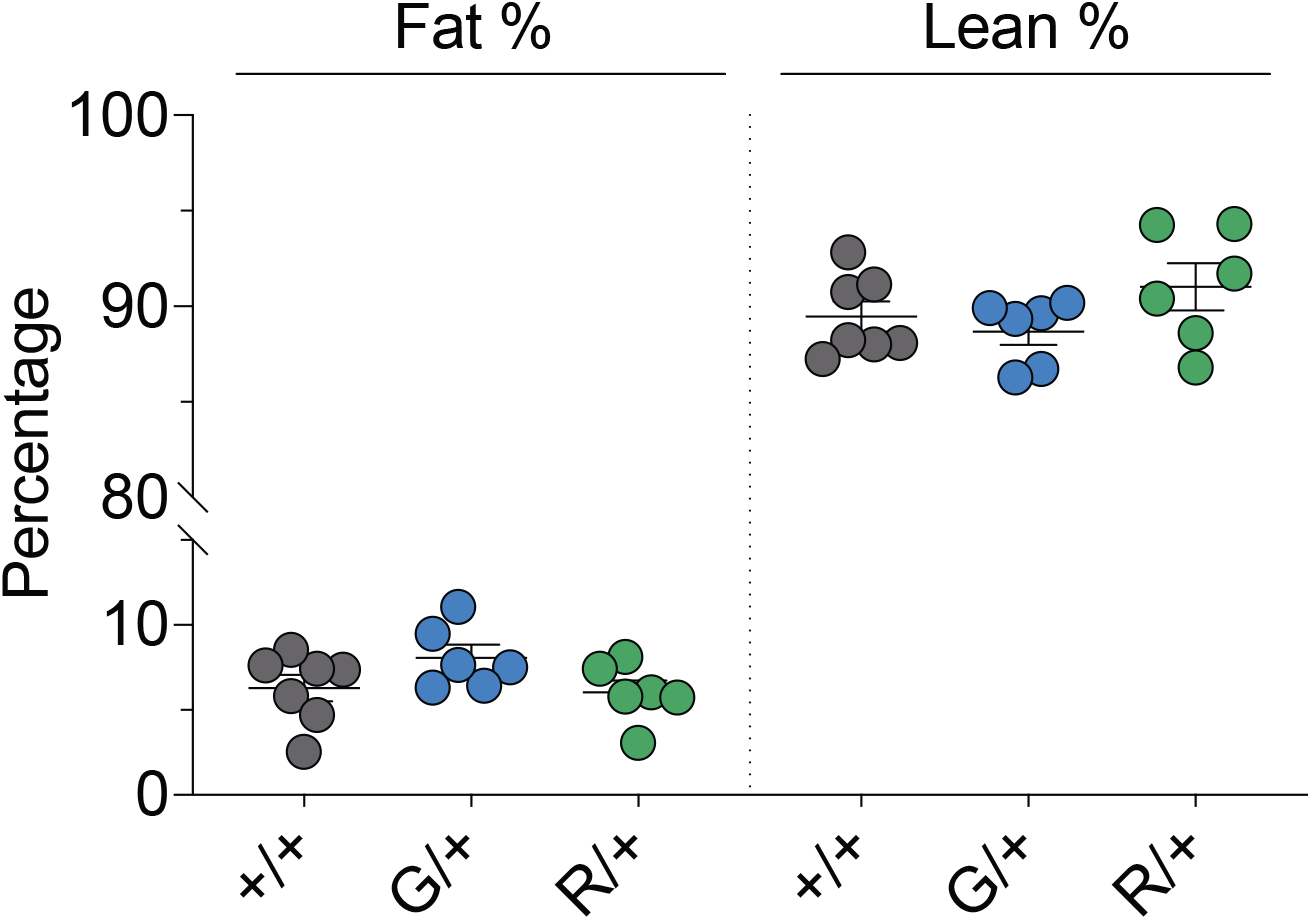
The reduced size of the heterozygous *Recql4* mutants is not a result of a change in fat to lean mass ratio. Echo-MRI analysis of fat and lean percentage at ten weeks of age from males *Recql4^+/+^, Recql4^G522Efs/+^,* and *Recql4^R347X/+^* mice.

**S4 Fig.**
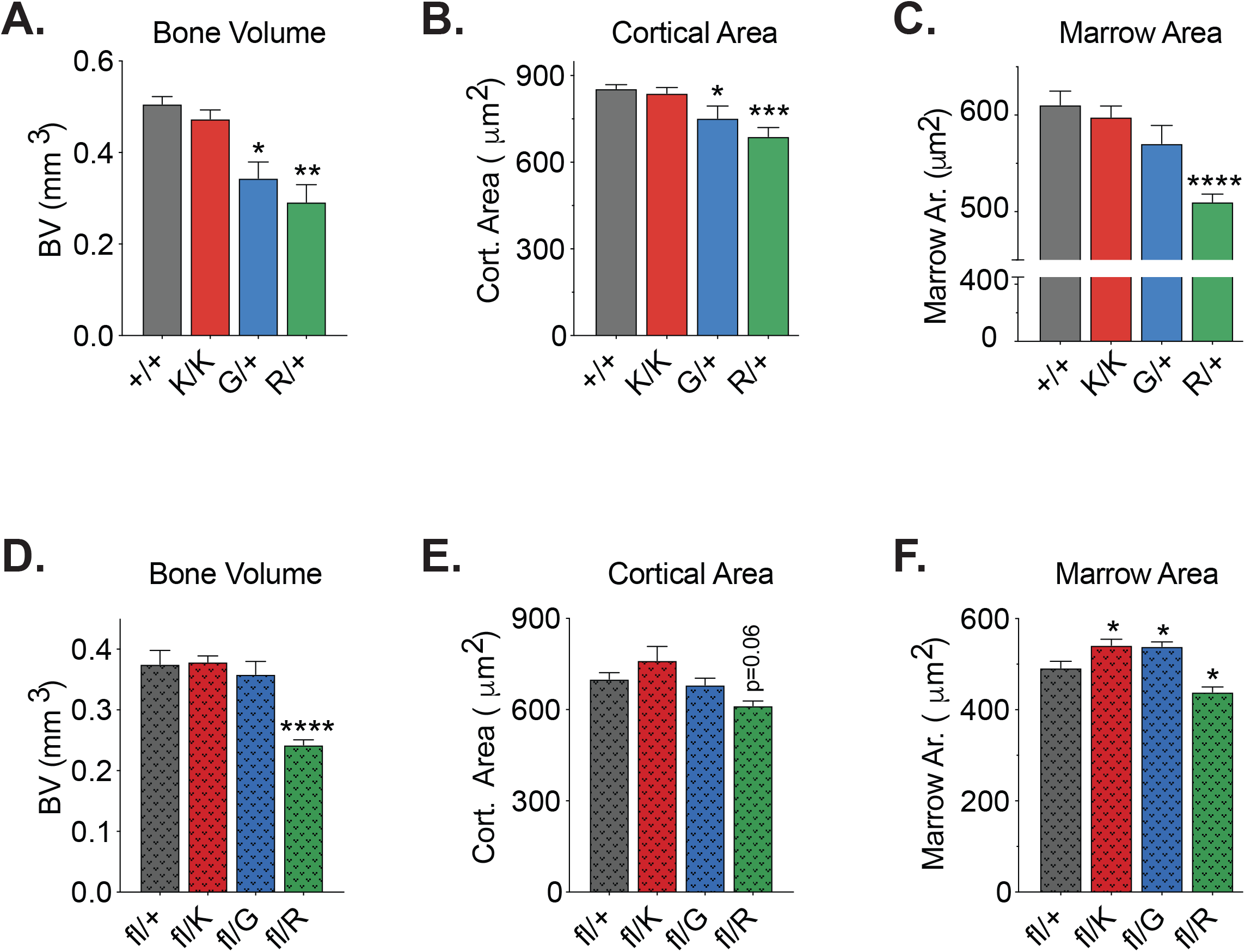
Micro-CT analysis. Micro-CT measurements of germline 10-week old males *Recql4^+/+^, Recql4^K525A/K525A^, Recql4^G522Efs/+^,* and *Recql4^R347X/+^* mice: (A) Bone volume. (B) Cortical area. (C) Marrow area. Micro-CT measurements of 10-week old males *Osx*-Cre *Recql4^fl/+^, Osx*-Cre *Recql4^fl/K525A^, Osx*-Cre *Recql4^fl/G522Efs^,* and *Osx*-Cre *Recql4^fl/R347X^* mice: (D) Bone volume. (E) Cortical area. (F) Marrow Area. Data expressed as mean ± SEM, Ordinary one-way ANOVA. *P<0.05; **P<0.01; ***P<0.001; ****P<0.0001; n≥6 per genotype. Experiments were independently executed on separate cohorts, with results pooled for presentation. K=K525A; G=G522Efs; R=R347X.

**S5 Fig.**
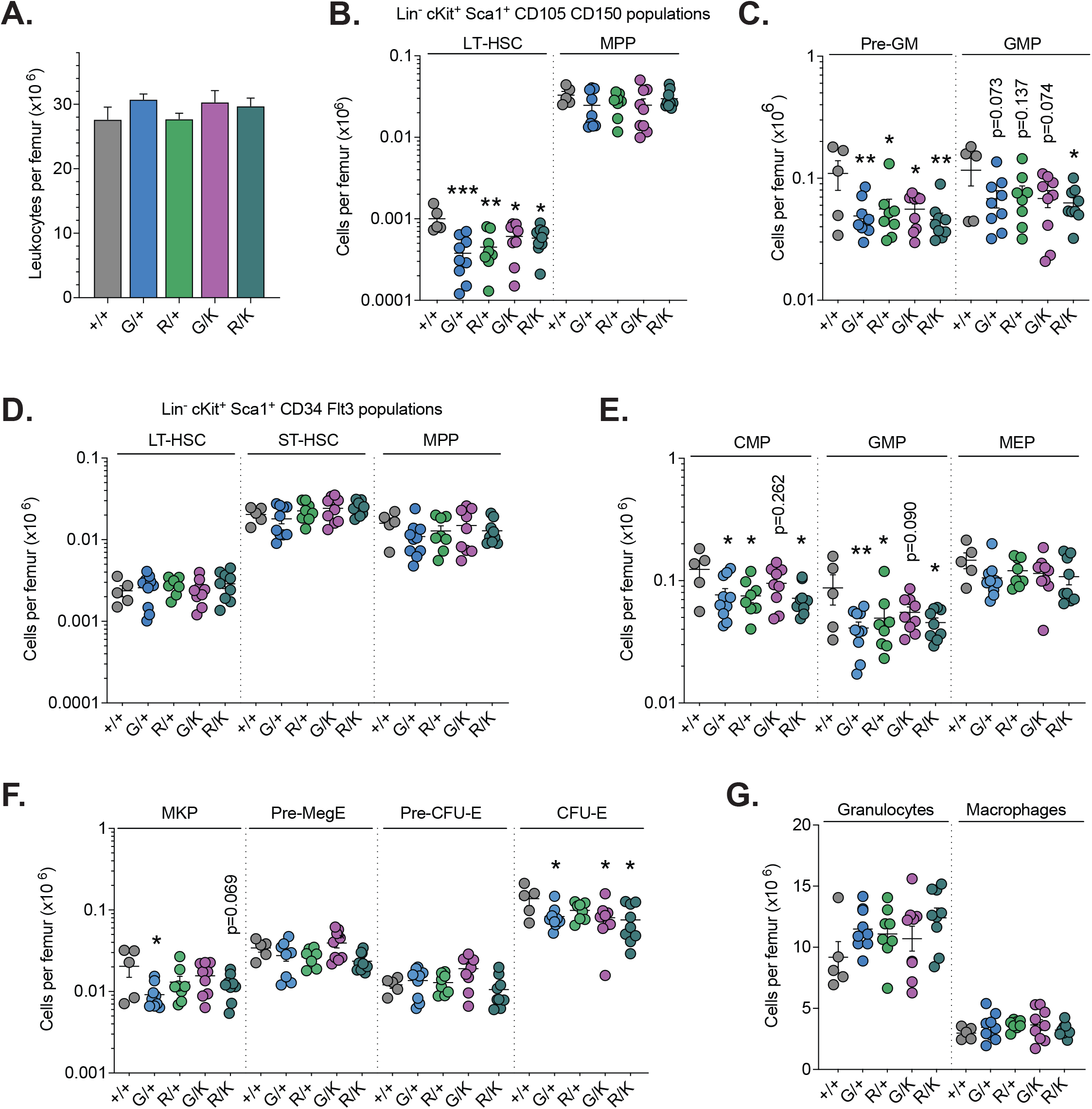
Heterozygous *Recql4* truncating mutations mildly affect myeloid progenitor populations. (A) Total leukocyte counts. (B) Hematopoietic stem cells (HSC) and primitive progenitors based on Lin-c-kit+Sca-1+CD105/CD150 staining per femur. (C) Pre-GM and GMP populations per femur. (D) HSC and primitive progenitors based on Lin-c-kit+Sca-1+CD34/Flt3 staining per femur. (E) Numbers of myelo-erythroid progenitors (MEP) per femur. (F) Erythroid (Pre-CFU-E, CFU-E) and megakaryocyte progenitor (MkP) frequency in the bone marrow and representative FACS plot. (G) Numbers of granulocytes and macrophages per femur. Data expressed as mean ± SEM, ordinary one-way ANOVA. *P<0.05; **P<0.01; ***P<0.001. Experiments were independently executed on separate cohorts, with results pooled for presentation. +/+=wild type, n=5; G=G522Efs, n=10; R=R347X, n=8; G/K=G522Efs/K525A, n=9; R/K=R347X/K525A, n=9.

**S6 Fig.**
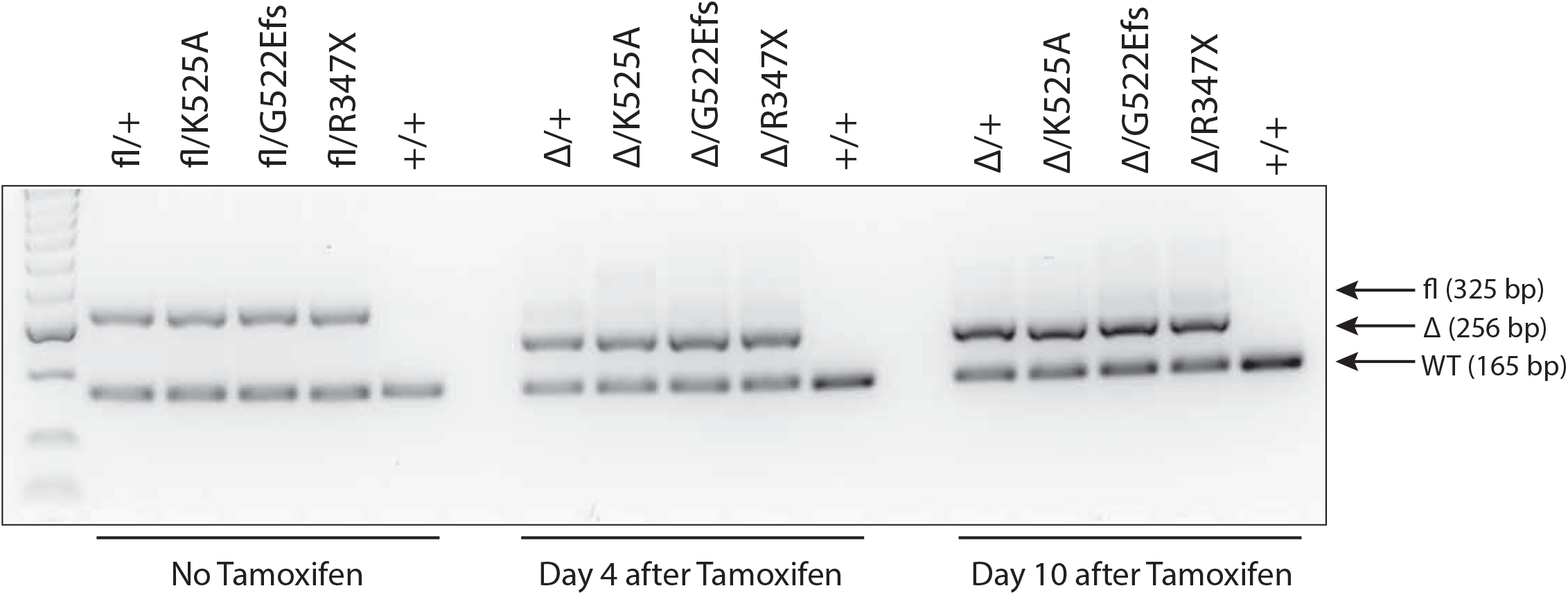
Effective deletion of *Recql4* floxed allele in murine mutants. Genomic PCR from Hoxb8 immortalized *R26*-CreER^T2^ *Recql4^fl/+^*, *R26*-CreER^T2^ *Recql4^f/lK525A^, R26*-CreER^T2^ *Recql4^fl/G522Efs^,* and *R26*-CreER^T2^ *Recql4^fl/R347X^* after addition of tamoxifen (400nM/mL) for 14 days demonstrating efficient gene deletion. True wild type (+/+) was used as control.

**S7 Fig.**
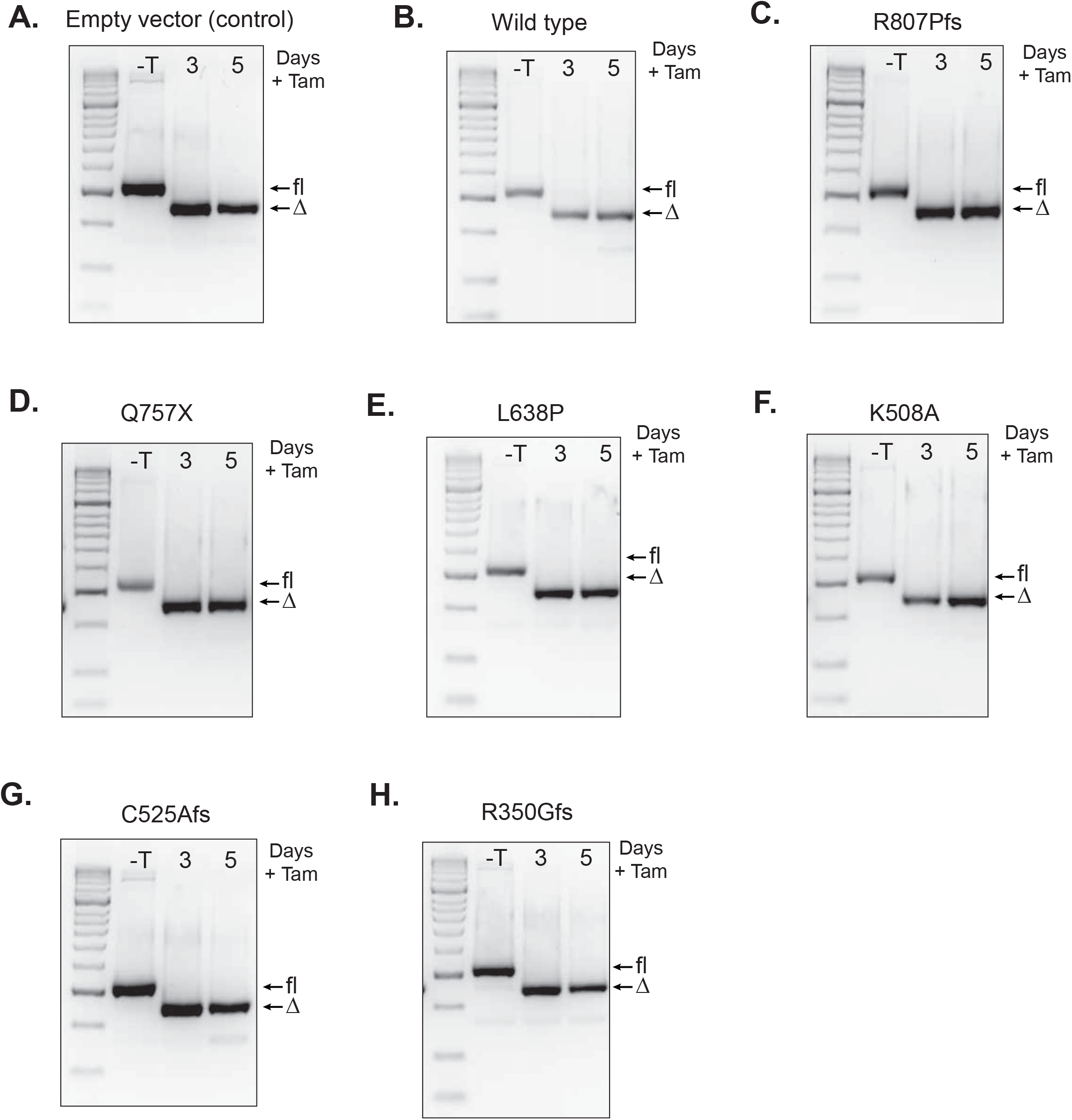
Effective deletion of *Recql4* floxed alleles in cells expressing human constructs. Genomic PCR from Hoxb8 immortalized *R26*-CreER^T2^ *Recql4^fl/f^* cells expressing: (A) empty vector (control); or EGFP fused (B) wild type RECQL4; (C) R807Pfs; (D) Q757X; (E) L638P; (F) K508A; (G) C525Afs; (H) R350Gfs. Efficient gene deletion was achieved after the addition of tamoxifen (400nM/mL) for the indicated number of days.

**S8 Fig.**
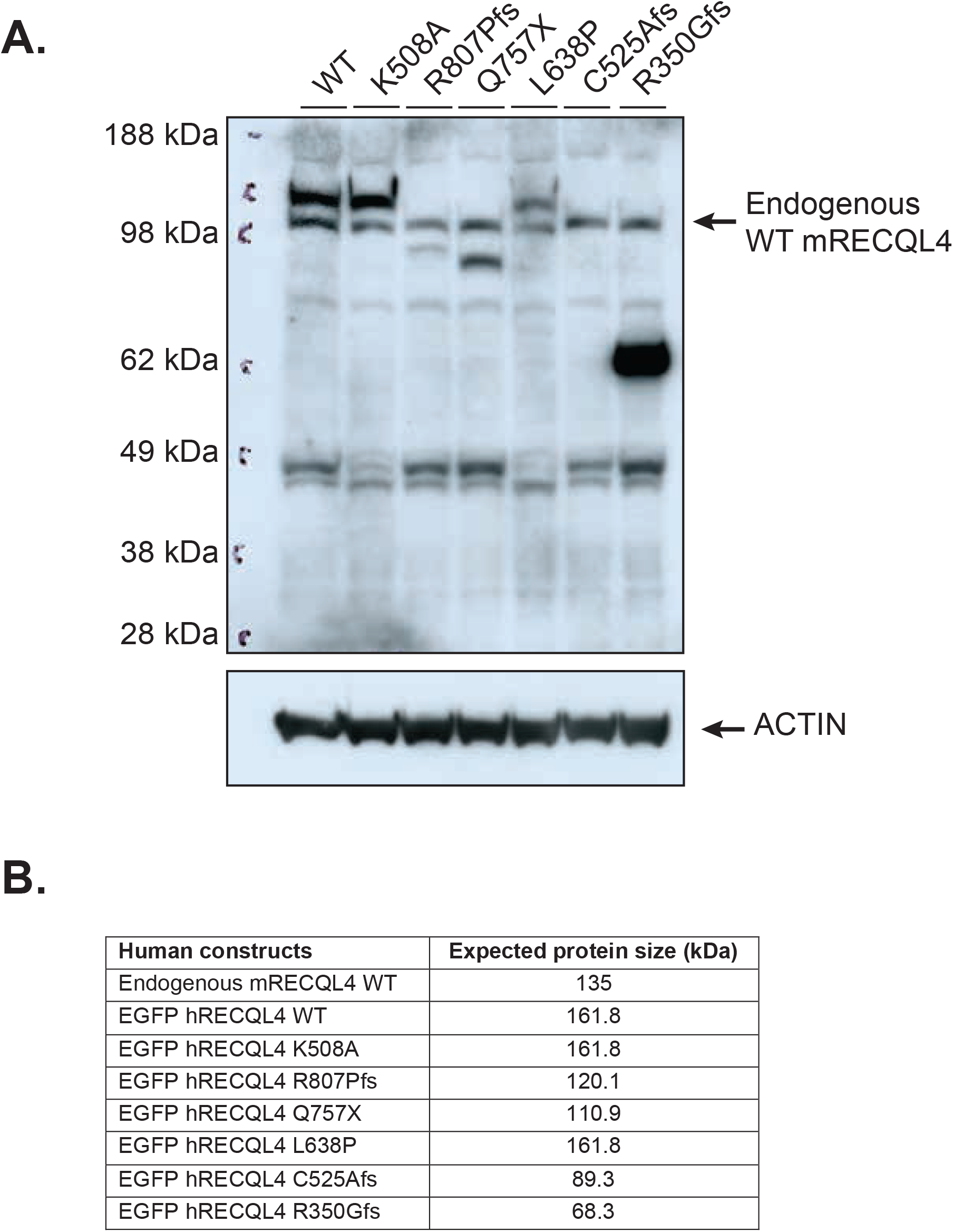
Protein expression of human constructs. (A) Western blot from Kusa4b10 cells expressing EGFP fused RECQL4 wild type, K508A, R807Pfs, Q757X, L638P, C525Afs, and R350Gfs. Cells probed with anti-human/mouse RECQL4 (clone 3B1; top). The same blot re-probed with anti-Actin (bottom). (B) Expected protein product sizes.

**S9 Fig.**
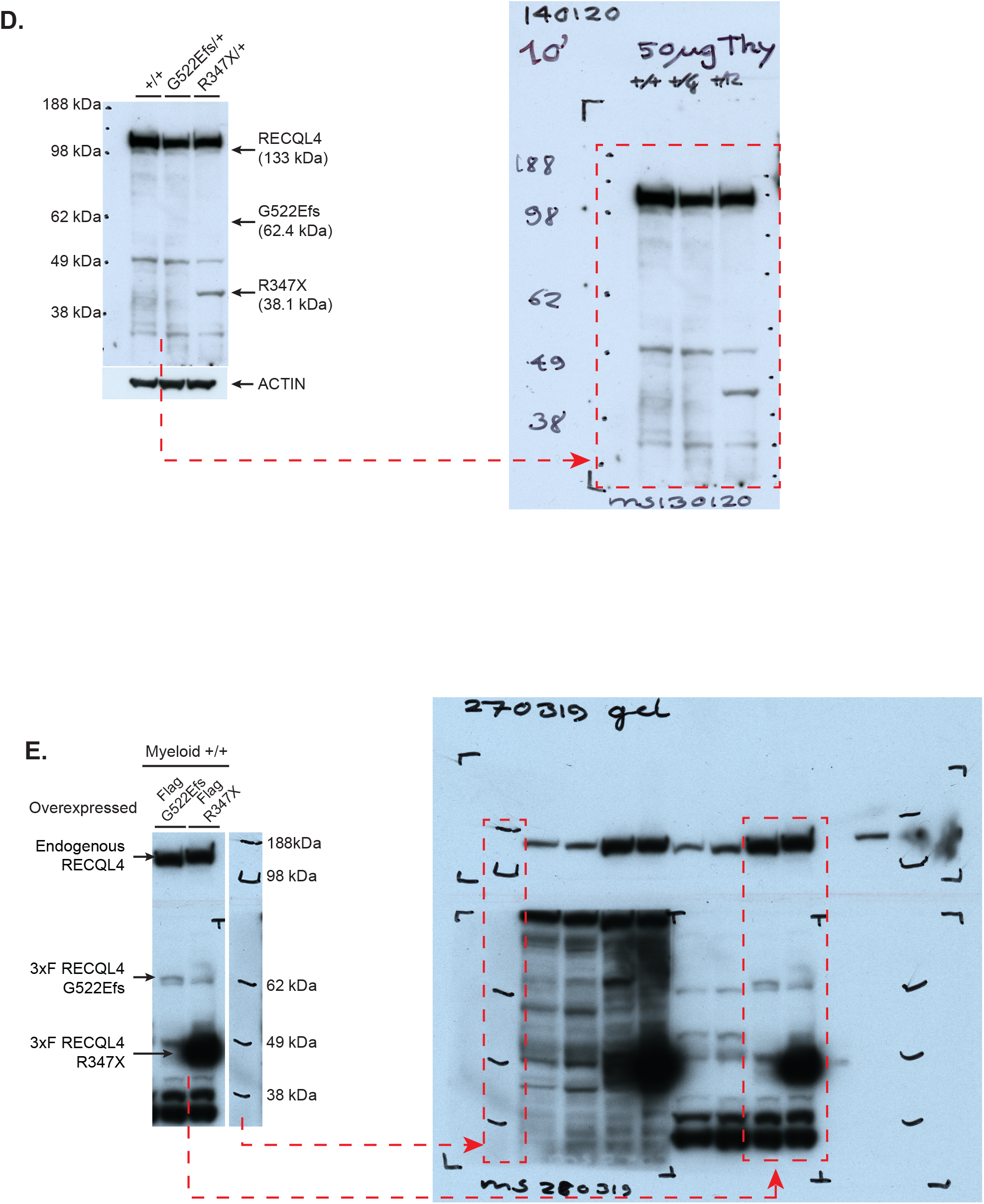
Uncropped westerns for Figure 1D and Figure 1E.

**S1 Table.** Tumors observed in mutant mice. List of tumors found during autopsy.

| **Tumor type** | **Recql4<sup>+/+</sup>** | **Recql4<sup>K525A/+</sup>** | **Recql4<sup>K525A/K525A</sup>** | **Recql4<sup>G522/K525A</sup>** | **Recql4<sup>R347X/K525A</sup>** |
| --- | --- | --- | --- | --- | --- |
| Intestinal tumor | 1 | 3 | 1 | 2 |  |
| Thymus tumor | 2 |  | 2 |  | 1 |
| Hepatic tumor | 2 | 3 | 2 | 2 | 2 |
| Lung tumor | 1 |  |  | 1 |  |
| Spleen tumor |  |  |  | 1 | 1 |
| Mammary gland tumour |  |  | 1 |  |  |

**S2 Table.** FACS Antibodies. List of antibodies used for flow cytometry in this study.

| **Antibody (clone)** | **Conjugate** | **Catalogue #** | **Company** |
| --- | --- | --- | --- |
| Ter119 (TER-119) | PE | 12-5921-83 | Life Technologies Australia Pty Ltd/Thermo Fisher |
| CD71 (RI7 217.1.4) | APC | 17-0711-82 | Life Technologies Australia Pty Ltd/Thermo Fisher |
| B220/CD45R (RA3-6B2) | APC | 17-0452-83 | Life Technologies Australia Pty Ltd/Thermo Fisher |
| IgM (II/41) | Biotin | 13-5790-82 | Life Technologies Australia Pty Ltd/Thermo Fisher |
| CD43 (S7) | PE | 553271 | BD Pharmingen |
| CD19 (1D3) | PerCP-Cy5.5 | 45-0193-82 | Life Technologies Australia Pty Ltd/Thermo Fisher |
| Mac1/CD11b (M1/70) | PE | 12-0112-83 | Life Technologies Australia Pty Ltd/Thermo Fisher |
| Gr-1/Ly6G (RB6-8C5) | PE-Cy7 | 25-5931-82 | Life Technologies Australia Pty Ltd/Thermo Fisher |
| F4/80 (BM8.1) | APC | 20-4801-U100 | Tonbo Biosciences |
| CD4 (RM4-5) | eFluor-450 | 48-0042-82 | Life Technologies Australia Pty Ltd/Thermo Fisher |
| CD8a/Ly-2 (53-6.7) | APC-eFluor 780 | 47-0081-82 | Life Technologies Australia Pty Ltd/Thermo Fisher |
| TCRb (H57-597) | PE | 12-5961-83 | Life Technologies Australia Pty Ltd/Thermo Fisher |
| CD25 (PC61.5) | PE | 12-0251-83 | Life Technologies Australia Pty Ltd/Thermo Fisher |
| CD44 (IM7) | APC | 17-0441-83 | Life Technologies Australia Pty Ltd/Thermo Fisher |
| Sca-1 (D7) | APC | 17-5981-82 | Life Technologies Australia Pty Ltd/Thermo Fisher |
| c-Kit/CD117 (2B8) | APC-eFluor-780 | 47-1171-82 | Life Technologies Australia Pty Ltd/Thermo Fisher |
| CD34 (RAM34) | eFluor-660 | 50-0341-82 | Life Technologies Australia Pty Ltd/Thermo Fisher |
| CD135/Flk2/Flt3 (A2F10) | PE | 12-1351-82 | Life Technologies Australia Pty Ltd/Thermo Fisher |
| FcγR/CD16/32 (93) | PerCP-Cy5.5 | 45-0161-82 | Life Technologies Australia Pty Ltd/Thermo Fisher |
| CD41 (MWReg30) | eFluor-450 | 48-0411-82 | Life Technologies Australia Pty Ltd/Thermo Fisher |
| CD105 (MJ7/18) | PE-Cy7 | 120409 | Biolegend |
| CD150 (TC15-12F12.2) | PE | 115904 | Biolegend |
| Streptavidin | BV 605 | 563260 | BD Pharmingen |

